# Gut microbial-mediated polyphenol metabolism is restrained by parasitic whipworm infection and associated with altered immune function in mice

**DOI:** 10.1101/2024.02.06.579078

**Authors:** Audrey Inge Schytz Andersen-Civil, Pankaj Arora, Ling Zhu, Laura J. Myhill, Nilay Büdeyri Gökgöz, Josue L. Castro-Mejia, Milla M. Leppä, Lars H. Hansen, Jacob Lessard-Lord, Juha-Pekka Salminen, Stig M. Thamsborg, Dennis Sandris Nielsen, Yves Desjardins, Andrew R. Williams

**Author notes:** Joint First Authors (Audrey Inge Schytz Andersen-Civil, Pankaj Arora).

## Abstract

Polyphenols are phytochemicals commonly found in plant-based diets which have demonstrated immunomodulatory and anti-inflammatory properties. However, the interplay between polyphenols and pathogens at mucosal barrier surfaces has not yet been elucidated in detail. Here, we show that proanthocyanidin (PAC) polyphenols interact with gut parasites to influence immune function and gut microbial-derived metabolites in mice. PAC intake inhibited mastocytosis during infection with the small intestinal roundworm *Heligmosomoides polygyrus*, and induced a type-1, interferon-driven mucosal immune response during infection with the large intestinal whipworm *Trichuris muris.* PAC also induced alterations in mesenteric lymph node T-cell populations that were dependent on infection model, with a Th2/Treg bias during *H. polygyrus* infection, and a Th1 bias during *T. muris* infection. In the absence of infection, PAC intake promoted the expansion of *Turicibacter* sp. within the gut microbiota, increased faecal short chain fatty acids, and enriched phenolic metabolites such as phenyl-γ-valerolactones in the caecum. However, these putatively beneficial effects were reduced in PAC-fed mice infected with *T. muris,* suggesting concomitant parasite infection can attenuate gut microbial-mediated PAC catabolism. Collectively, our results suggest an inter-relationship between a phytonutrient and infection, whereby PAC may augment parasite-induced inflammation (most prominently with the caecum dwelling *T. muris*), and infection may abrogate the beneficial effects of health-promoting phytochemicals.

## Introduction

Interest in the application of dietary phytonutrients as immunomodulatory and health-promoting dietary substances has been greatly increasing, due to rising rates of chronic autoimmune pathologies and antimicrobial drug resistance^1^. Proanthocyanidins (PAC) are plant-derived polyphenols, which are present in varying concentrations among commonly consumed foods, particularly grapes, berries and nuts^2^. Several studies have reported that PAC exhibit immunomodulatory properties, of which a common denominator is the downregulation of oxidative stress and aberrant inflammatory responses^3–7^. This may derive from direct modulation of antioxidant and inflammatory responses in gut epithelial and immune cells, as well as prebiotic effects whereby commensal gut bacteria metabolize PAC into phenolic metabolites that can improve gut barrier function^8–10^, or that are absorbed and subsequently exert systemic anti-inflammatory effects ^11–14^.

The intestinal tract is continuously challenged by potential harmful stimuli, while balancing the proportion of commensal and opportunistic bacteria^15, 16^. Therefore, the gut is one of the most immunologically active organs in the body, and the immune and inflammatory tone in the gut may impact not only responses to enteric pathogens but also inflammation and metabolism in extraintestinal tissues. The mucosal immune system contains discrete components that are specialized for combating specific infections, namely pro-inflammatory or type-1 responses against intracellular viruses and bacteria, and type-2 responses against multicellular parasitic worms (helminths). Dysregulation of these immune cell subsets can result in chronic disease such as Crohn’s disease, which is an overactive type-1 response, or food allergies that are driven by overactive type-2 responses^17, 18^.

Helminths are one of the most widespread pathogens of humans and animals, with around 1 billion people globally infected by different gastrointestinal-dwelling worms^19^. Further, these parasites are also ubiquitous in farmed livestock^20^. Under natural conditions, individuals are infected repeatedly with low-dose helminth infections throughout their lifetime, thus allowing the parasite to establish chronic, non-resolving infections^21^. When protective immunity to helminths develops, it is accompanied by type-2 immune mechanisms such as Th2 cells that produce IL-4 and IL-13, typically also accompanied by a strong T-regulatory component^22^. Murine infection models have established a clear paradigm that whilst enhancing the type-2 response increases resistance to helminths, strengthening of the type-1 response results in susceptibility to chronic infection^21, 23–25^.

In contrast to the well-known effects of PAC on improving health during chronic metabolic diseases, it is not yet understood in detail how they impact immune function during pathogen infections. PAC-rich diets have shown modulatory effects on the immune response to helminth infections in livestock (sheep and pigs), including a boosting of infection-induced γδ T-cells, eosinophils, and antibodies^26, 27^. However, PAC-enriched diets have also been shown to exacerbate infection with the extracellular bacterium *Citrobacter rodentium* infection in mice, which may be due to PAC-induced changes in the GM or mucosal immune function^28^. Thus, the effects of PAC on enteric infections may vary depending on factors such as host species or basal GM. So far, no studies have investigated the influence of PAC on helminth infection in mouse models fed purified, semi-synthetic diets (SSD) with a defined chemical composition. This may represent a useful system to examine the interactions between PAC and gut pathogens, and to assess if PAC intake can modulate the polarization of the immune system towards either a type-1 or type-2 immune response in natural models of parasite infection.

Here, we explored if the addition of PAC to purified, open-source murine diets can alter the immune response during enteric parasitic infection. We utilized two parasite systems, the small intestinal *Heligmosomoides polygyrus* and the large intestinal *Trichuris muris*, which can be used to effectively model the infection dynamics typically seen in natural infections in humans and animals^22, 29^. We show that in both infection models PAC induces a modulation of the immune response induced by infection. Moreover, infection tended to modulate PAC-induced changes in the gut microbiome. Importantly, whilst PAC intake increased the abundance of microbial-derived metabolites positively associated with gut health, concurrent *T. muris* infection impaired production of such metabolites. Thus, our results point to an interaction between enteric parasite infection and dietary phytonutrients that may, in some contexts, negatively affect host health.

## Results

### Proanthocyanidins alter expression of immune-related genes and reduce mastocytosis during small intestinal nematode infection

To study if PAC can modulate mucosal immune responses to an enteric helminth infection, we first examined the effects of PAC intake on the small intestinal mucosal immune response to *H. polygyrus*. Mice were fed SSD (Supplementary Table 1) and were gavaged every 2^nd^ day with 200 mg/kg purified PAC derived from grape pomace dissolved in water, or water only, for 4 weeks. Halfway through the experiment, mice in each treatment group were infected with *H. polygyrus* and euthanized 14 days post-infection (p.i.,), i.e., 28 days after initiation of PAC administration, or remained as uninfected controls (**Figure 1A)**. RNA sequencing of duodenum tissues from *H. polygyrus-*infected mice without PAC intake showed a strong upregulation of many genes involved in type-2 immunity, relative to uninfected control mice (**Figure 1B**; Supplementary File 1). These included genes involved in mast cell responses, including mast cell proteases (*Mcpt1, Mcpt2*), chymases (*Cma2*) and subunits of the high-affinity IgE receptor (*Ms4a2*). Furthermore, innate defense molecules (e.g., *Ang4*) were upregulated (**Figure 1B**). Downregulated genes were mainly involved in nutrient uptake and metabolism, consistent with the marked changes in gut function that accompany *H. polygyrus* infections^30^. Transcriptional pathways enriched by *H. polygyrus* infection were mainly related to mastocytosis, whilst suppressed pathways were connected to metabolic processes such as electron transfer, ATP synthesis and cellular biotransformation (**Figure 1C**). Thus, *H. polygyrus* infection induced a strong modulatory effect on the intestinal environment with marked upregulation of type-2 immune mechanisms, and a suppression of cellular metabolism that may relate to mucosal injury and necrosis induced by nematode invasion of the host tissue.

**Figure 1.**
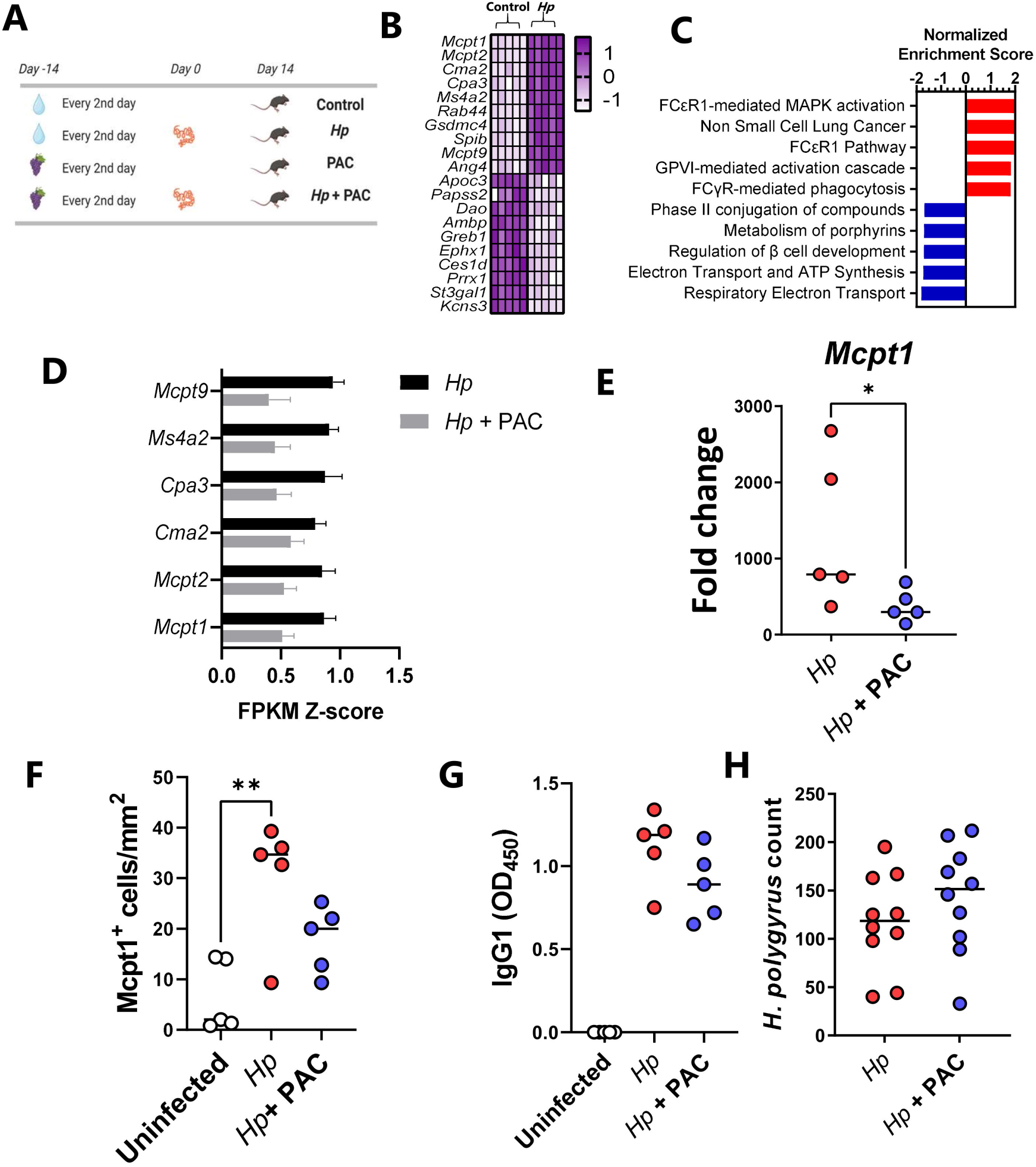
Proanthocyanidins downregulate immune-related genes and affect immune cell hyperplasia in duodenum tissues of *Heligmosomoides polygyrus*-infected mice with no effect on worm burdens. **A)** Schematic of experimental design. Mice in proanthocyanidin (PAC)-treated groups received 200 mg/kg PAC derived from grape pomace every 2^nd^ day, whilst control mice received water. Mice infected with *H. polygyrus* (*Hp*) were inoculated with 200 larvae on day 0. **B)** Top ten up- and down-regulated genes (Z-scores) in the duodenum of *H. polygyrus*-infected mice relative to uninfected control mice, identified by RNA-sequencing (for all genes adjusted *p* value <0.05 by DeSeq2). **C)** Top 5 up- and down-regulated gene pathways (*q* value <0.05) in duodenal tissue identified by gene set enrichment analysis in mice infected with *H. polygyrus*, relative to uninfected mice. **D)** Specific expression of mast cell-related genes in the duodenum of *H. polygyrus*-infected mice and PAC-dosed *H. polygyrus*-infected mice, relative to uninfected control mice. **E)** Expression of *Mcpt1* as measured by qPCR. Fold changes are relative to uninfected, control mice. **F)** Enumeration of Mcpt1-positive mast cells in the duodenum. **G)** *H. polygyrus*-specific serum IgG1 levels in serum **H)** Worm burdens in *H. polygyrus*-infected mice with and without PAC treatment. **A)-G),** n=5 per treatment group from a single experiment, **H)** n=10 per group, pooled from two independent experiments. For all panels medians are shown. *p < 0.05, **p < 0.01, by Mann-Whitney tests or Kruskal-Wallis tests with Dunn’s post-hoc testing.

In contrast, PAC intake in uninfected mice did not substantially change the duodenal transcriptome, with no significantly regulated genes following multiple correction testing (Supplementary File 2). PAC do therefore not seem to have a strong effect at this tissue site, at least in parasite-naïve mice. To explore if PAC intake impacted on host responses during *H. polygyrus* infection, we next compared duodenal transcriptomic responses of *H. polygyrus* - infected mice receiving either PAC or water alone. Overall, PAC had only minor effects on the transcriptional response to *H. polygyrus* infection. However, we noted suppression of several genes by PAC that were related to mast cell or immunoglobulin signaling, such as *Mcpt1* and *Ms4a2*, albeit again not significantly so after correction for multiple testing (Supplementary Figure 1; Supplementary File 3). Indeed, expression of the top mast cell related genes identified as being upregulated by *H. polygyrus* in Figure 1A was noticeably lower in infected mice administered PAC, relative to infected controls (**Figure 1D**). Given the lack of statistical significance from RNA-seq analysis, we verified the suppression of the type-2 response in several ways. qPCR confirmed that infection-induced upregulation of *Mcpt1* was significantly attenuated by PAC, and Mcpt1^+^ mast cells in the intestinal mucosa were significantly elevated by infection in infected control-fed mice, but not in infected mice administered PAC (**Figure 1E-F**). However, we observed no significant effect of PAC on *H. polygyrus*-specific IgG1 levels in serum or worm burdens at day 14 p.i., (**Figure 1G-H)**. Thus, in this model dietary PAC supplementation did not significantly enhance parasite-specific immunity, but rather tended to restrain type-2 immune responses induced by *H. polygyrus*, although worm counts remained unchanged.

### Proanthocyanidins modulate caecal transcriptomic responses and alter *Trichuris muris* infection dynamics

To further investigate the effect of PAC intake on helminth-specific immune function, we utilized a second infection model, the caecal dwelling whipworm *T. muris*, that also closely resembles the chronic infection and inflammation that may occur in humans and livestock during natural exposure. Low doses of *T. muris* eggs induce both type-1 and type-2 immune effector mechanisms, with the type-1 bias allowing worm persistence and the development of colitis-like pathology^29^. Furthermore, the location of *T. muris* in the caecum, the main site of fermentation of diet-derived plant material in mice, may make this model particularly relevant for assessing interactions with dietary components, given that PAC consumption has been previously shown to modulate gene expression and gut barrier activity in the caecum and colon^31, 32^.

Mice trickle infected with *T. muris* (3 doses of 20 eggs over 35 days), and uninfected controls, received oral administration of purified PAC from grape seed extract (300 mg/kg) 14 days prior to, and throughout, the infection period (**Figure 2A**). Interestingly, PAC intake decreased body weight gains relative to mice consuming only SSD (Supplementary Figure 2). Using RNA-sequencing, we first characterized the transcriptional response in caecum tissue to *T. muris* infection alone. Relative to uninfected controls, *T. muris*-infected mice had altered expression of more than 2000 genes (adjusted *p* <0.05; Fold change ≥2). Similar to *H. polygyrus* infection, mast-cell related genes such as *Mcpt1* were strongly upregulated but were accompanied by pronounced upregulation of *Gzma*, *Ifng*, and *Nos2*, suggesting a mixed type-1/type-2 response (**Figure 2B**; Supplementary File 4). Consistent with this, gene pathways indicative of pro-inflammatory response (NK cell signaling, CD28 signaling) were strongly enriched, whilst metabolic pathways connected to hormone and peroxisome proliferator-activated receptor (PPAR) signaling were amongst the most suppressed (**Figure 2C**). Thus, *T. muris* trickle infection induced significant caecal inflammation and disruption of metabolic homeostasis in the gut.

**Figure 2.**
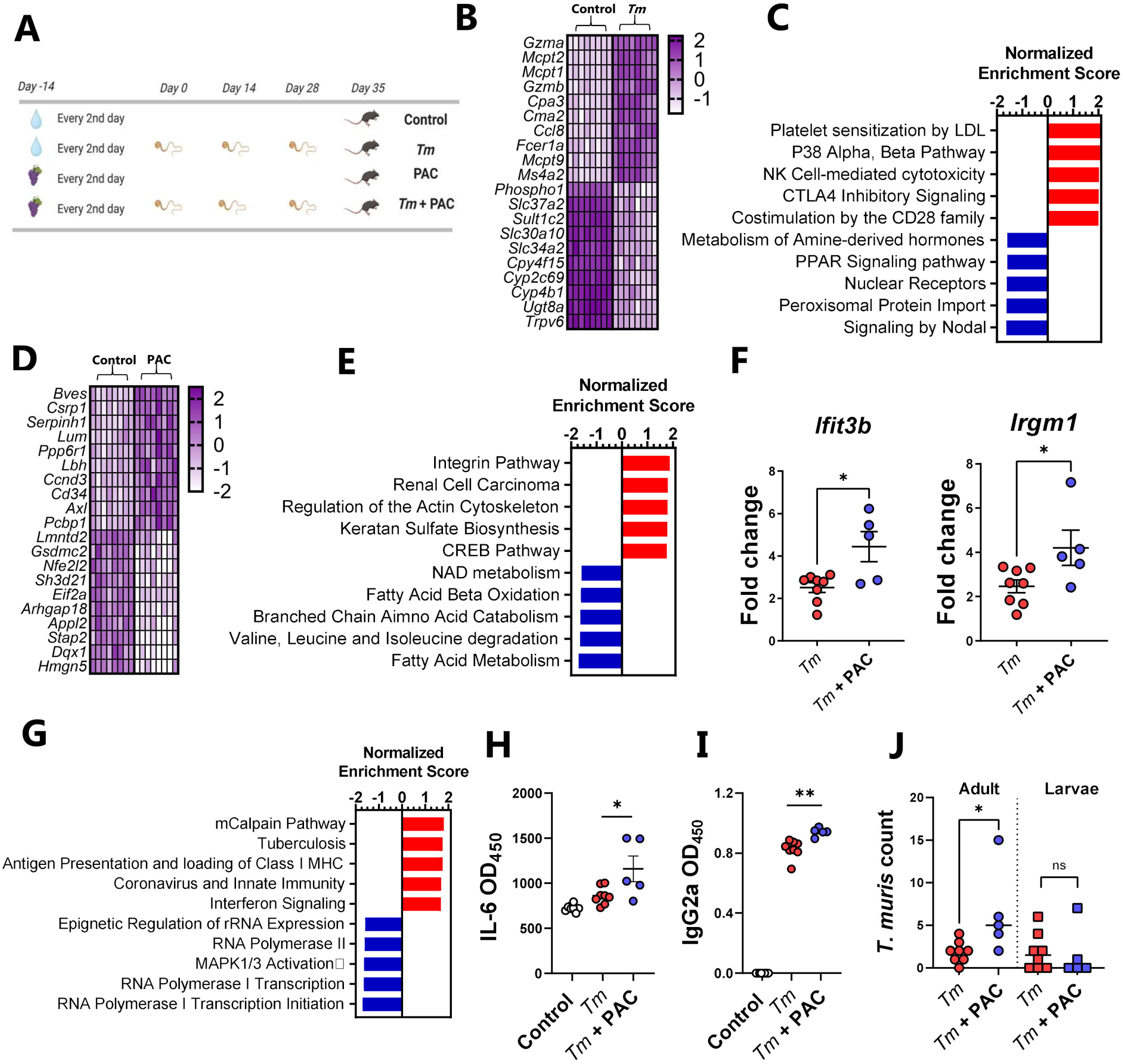
Proanthocyanidins modulate immune responses in *Trichuris muris*-infected mice. **A)** Schematic of experimental design. Mice in proanthocyanidin (PAC)-treated groups received 300 mg/kg PAC derived from grape seeds every 2^nd^ day, whilst control mice received water. Mice infected with *Trichuris muris* (*Tm*) were inoculated with 20 eggs on day 0, 14 and 28. **B)** Top ten up- and down-regulated genes (Z-scores) in the caecum of *T. muris*-infected mice relative to uninfected control mice, identified by RNA-sequencing (for all genes adjusted *p* value <0.05 by DeSeq2). **C)** Top 5 up- and down-regulated gene pathways (*q* value <0.05) in caecal tissue identified by gene set enrichment analysis in mice infected with *T. muris,* relative to uninfected mice. **D)** Top ten up- and down-regulated genes (Z-scores) in the caecum of PAC-dosed mice relative to water-dosed control mice, identified by RNA-sequencing (for all genes adjusted *p* value <0.05 by DeSeq2). **E)** Top 5 up- and down-regulated gene pathways (*q* value <0.05) in caecal tissue identified by gene set enrichment analysis in PAC-dosed mice relative to water-dosed control mice. **F)** Top 5 up- and down-regulated gene pathways (*q* value <0.05) in caecal tissue identified by gene set enrichment analysis in PAC-dosed mice infected with *T. muris,* relative to mice infected with *T. muris* alone. **G)** Expression of *Ifit3b* and *Irgm1* measured by qPCR. Fold changes are relative to uninfected, control mice. **H)** IL-6 induced by *T. muris* antigens in lymphocytes isolated from mesenteric lymph nodes in *T. muris*-infected mice, with or without PAC dosing. **I)** *T. muris*-specific serum IgG2a levels in serum. **J)** Adult and larval worm burdens in *T. muris*-infected mice For all panels, n=5-8 per group, pooled from two independent experiments. **G) – H) –** Shown are means ± S.E.M. *p < 0.05 by un-paired t test. **I) – J)** – Shown are median values. *p < 0.05, **p < 0.01 by Mann-Whitney test.

As a first step in determining whether PAC intake could modulate this response, we first carried out a similar RNA-seq approach to determine transcriptional responses in caecal tissue of uninfected mice fed PAC to establish a baseline profile of the effect of PAC administration. Most of the upregulated genes resulting from PAC intake (e.g. *Bves, Csrp1, Cd34*) were related to cell adhesion and differentiation, which was supported by the enrichment of gene pathways related to integrins, cell proliferation, and extracellular matrix remodeling (Actin, Keratan sulfate activity), consistent with known functions of PAC in stimulating epithelial cell growth^33^ (**Figure 2D-E**; Supplementary File 5). Interestingly, PAC intake was associated with down-regulation of *Nfe2l2,* encoding the transcription factor Nrf2, supporting a role in modulation of oxidative stress responses, as well as *Gsdmc2*, encoding the epithelial cell protein gasdermin-2 which may be involved with cell lysis and pyroptosis^34^. Downregulated gene pathways were mainly related to metabolic function such as fatty acid metabolism, coherent with known roles of PAC in regulating lipid and bile acid metabolism^35^. Thus, PAC intake, in the absence of infection, was associated with modulation of transcriptional pathways putatively associated with tissue remodeling and nutrient metabolism, suggesting that PAC has significant effects on the local mucosal environment in the large intestine.

Next, caecal tissues harvested from *T. muris-*infected mice administered PAC were compared to caecal tissues from mice infected with *T. muris* without PAC treatment. Notably, despite the differences in tissue type and infection dynamics we observed a similar picture to that observed with *H. polygyrus*. Specifically, only minor gene expression changes were induced by PAC, relative to water-dosed controls, with no genes being significantly changed after correction for multiple testing (Supplementary Figure 3; Supplementary File 6). However, when considering genes upregulated by PAC with unadjusted *p* values of <0.005, it was again notable that some of these (e.g. *Ifit3b* and *Irgm1*) were related to a shift from a type-2 to a type-1 immune environment in the infected tissue (Supplementary Figure 3). Thus, we explored in greater detail if PAC modulated the mucosal response to *T. muris*. First, we conducted gene-set enrichment analysis, which revealed significant (*q* <0.05) up-regulation of several pathways related to either interferon production, or immune responses towards viral or bacterial pathogens, in PAC-dosed mice during *T. muris* infection (**Figure 2F**). Next, we confirmed by qPCR that expression of *Ifit3b* and *Irgm1*, both encoding proteins downstream of IFNγ production, was significantly increased in caecal tissue of *T. muris*-infected mice fed PAC, relative to *T. muris* alone (**Figure 2G)**. We then analyzed serum antibodies and cytokine production from mesenteric lymph node (MLN) cells. Infected mice fed PAC had significantly increased serum levels of *T. muris-*specific IgG2a, a marker of type-1 responses and parasite chronicity^29^ (**Figure 2H)**. Whilst MLN cytokine secretion induced by *T. muris* antigen was largely similar between both groups (Supplementary Figure 4), we noted significantly higher IL-6 production in infected mice fed PAC (**Figure 2I**). Finally, adult worm burdens (but not larval, or total, worm burdens), were significantly higher in PAC-treated mice (**Figure 2J),** suggesting a modulation of parasite population dynamics as a result of the altered immune function. Taken together, these data indicate a significant enhancing effect of PAC on the type-1 polarized response elicited by parasitic infection, and collectively suggest that PAC intake during whipworm infection regulates host susceptibility to adult worms. Thus, despite PAC intake having seemingly beneficial effects on the mucosal barrier in uninfected mice, PAC intake during helminth infection was associated with an immune polarization which hampered the expression of protective immune mechanisms.

### Proanthocyanidins alter T cell populations in mesenteric lymph nodes in helminth-infected mice

The balance between differentiated intestinal T-cells during helminth infection may play a key role in determining the outcome of infection^36^. To explore if the transcriptomic changes observed in mucosal gut tissues were accompanied by changes in T-cell populations, MLN were isolated from *H. polygyrus*-infected mice at day 14 p.i. and from *T. muris*-infected mice at day 35 p.i. to assess how these were altered by infection and/or PAC supplementation. Interestingly, PAC intake significantly increased the total number of cells in the MLN of both infection models, relative to infected mice which were not administered PAC (**Figure 3A)**. PAC had no effect on proportions of Th1 (Tbet^+^), Th2 (GATA3^+^) and Foxp3^+^ T-helper cells within the T-helper cell population (TCRβ^+^CD4^+^) in uninfected mice (data not shown). In *H. polygyrus* -infected mice, proportions of Th1 cells were not affected by PAC, but, interestingly, the proportion of Th2 cells was significantly increased (**Figure 3B**), in contrast to the mast cell data overserved previously. However, we also noted a strong tendency for the proportion of Treg cells to also be increased by PAC (**Figure 3B**). In contrast, in *T. muris-*infected mice, PAC intake tended to increase proportions of Th1 cells in the MLN, and decrease Th2 or Foxp3^+^ cells, although no significant differences were observed (**Figure 3C)**. Thus, PAC supplementation had a stimulatory effect on lymphocyte proliferation in the MLN during helminth infection. However, the relative expansion of different T-cell subsets was highly dependent on infection model, with a seemingly opposed effect on the T-cell balance in *H. polygyrus* and *T. muris* infections.

**Figure 3.**
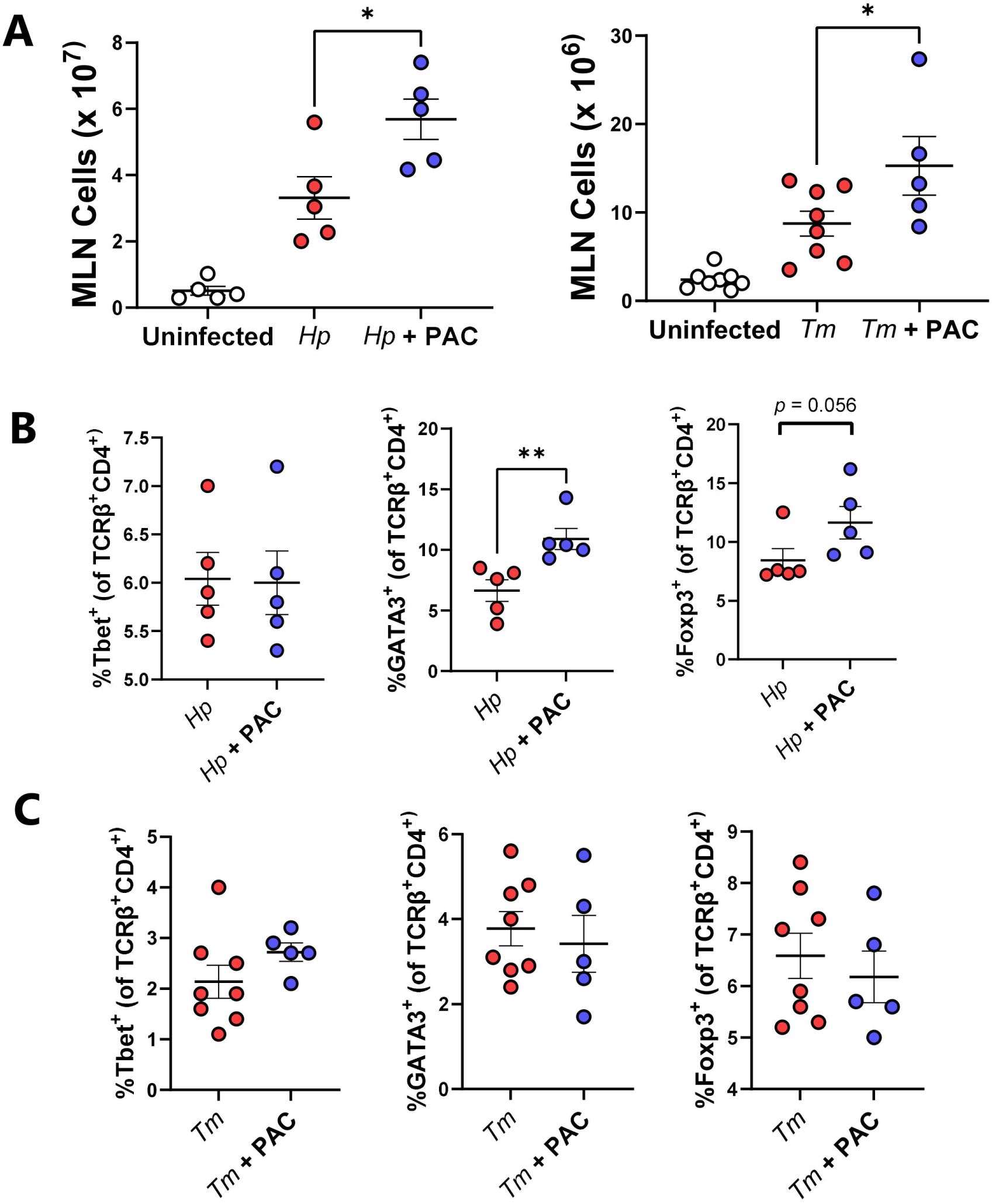
Proanthocyanidins alter T cell populations in the mesenteric lymph nodes in helminth-infected mice. Impact of proanthocyanidins (PAC) on the total number of cells (**A**), and proportions of TCRβ^+^ CD4^+^T-bet^+^, TCRβ^+^ CD4^+^GATA3^+^ and TCRβ^+^ CD4^+^Foxp3^+^ T-cells in the mesenteric lymph nodes (MLN) of *Heligmosomoides polygyrus* (*Hp*; **B**) and and *Trichuris muris* (*Tm*; **C**) infected mice. **B)** n=5 per treatment group from a single experiment. **C)** n=5-8 per group, pooled from two independent experiments. Shown are either means ± S.E.M. or median values. * *p*<0.05 by unpaired t-test. Gating strategy is shown in Supplementary Figure 7.

### Proanthocyanidins modulate immune responses during *Trichuris muris* infection more strongly with semi-synthetic diets than with chow

To explore if altered immune function during *T. muris* infection was dependent on the dietary matrix that accompanied PAC administration, we investigated the effect of basal diet composition on host responses to exogenous PAC supplementation. Previous studies have found that inclusion of plant-derived oligosaccharides in SSD, but not unrefined mouse chow, can exacerbate colitis and inflammation^37^, suggesting that basal diet may play a role in dictating how phytochemicals modulate gut inflammation. Mouse chow is a crude mixture containing many components including lignins, non-starch polysaccharides, and isoflavones. In contrast, SSD are essentially free of phytonutrients, containing mainly purified casein, starch, sucrose and cellulose (Supplementary Table 1). We thus hypothesized that the immunomodulatory effects of PAC would be more pronounced when administered together with SSD, than with mouse chow. Mice were trickle-infected with *T. muris* as above, and fed either chow or SSD, with or without 300 mg/kg purified PAC from grape seed extract every 2^nd^ day. In line with our previous results, mice dosed with PAC gained less weight, and this was most pronounced in the SSD-fed mice (**Figure 4A**). The pattern of worm burdens was similar to what we previously observed, with slightly higher adult worm burdens in PAC-dosed mice, although no significant differences were found. Strikingly, consistent with recent observations^38^, basal diet composition had a marked effect on *T. muris* burdens with a substantial increase in both adults and larvae in chow-fed mice, compared to SSD-fed mice (**Figure 4B-C**). There was also a significant increase in MLN cells in chow-fed mice compared to SSD-fed mice (**Figure 4D**). PAC administration did not significantly change the proportions of Th1 cells in the MLN compared to controls, but did reduce the proportion of Th2 cells, most prominently in the SSD-fed mice (p=0.06 for interaction between diet and infection; **Figure 4E-F)**. Thus, there was a significantly skewed Th1:Th2 ratio in SSD-fed mice (*p* < 0.05 for interaction between basal diet and PAC intake; **Figure 4G**). IgG2a levels were also markedly affected by basal diet, with higher levels in chow-fed mice, correlating with worm burden, with no significant effect of PAC (**Figure 4H**). Finally, we quantified *T. muris* antigen-induced IL-6 secretion in MLN cells and found increased levels in PAC-dosed mice, but again only when fed SSD and not chow (*p* <0.05 for interaction between basal diet and PAC intake; **Figure 4I**). Collectively, these data demonstrate that basal diet composition has a substantial effect on worm burden and immune responses, and that modulation of T-cell responses and IL-6 production in MLN during *T. muris* infection is more strongly influenced by PAC when mice are fed SSD than chow.

**Figure 4.**
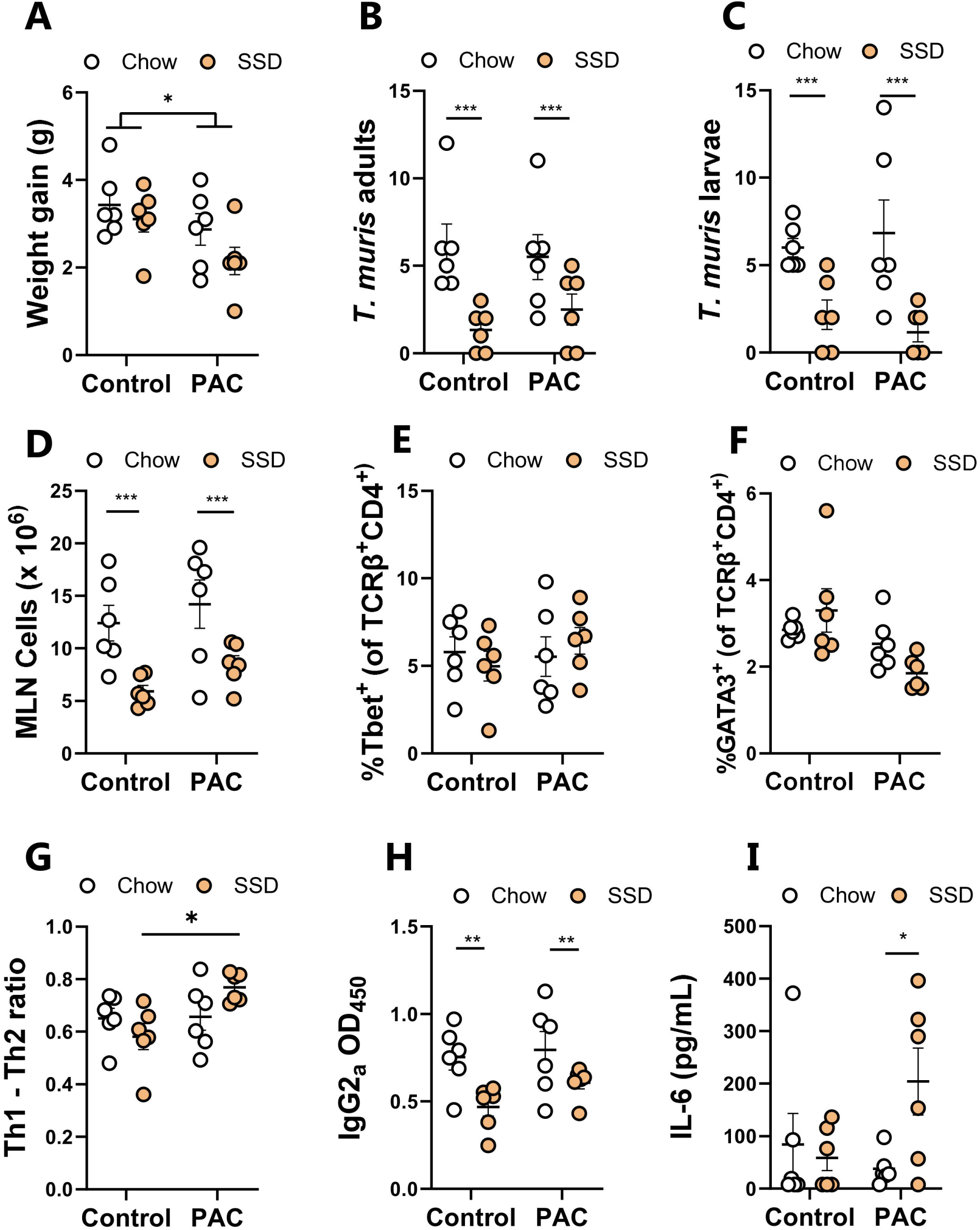
Effects of proanthocyanidins during *Trichuris muris* infection in mice fed either chow or semi-synthetic diets. **A)** Weight gain over the course of the 49 day experiment in mice fed either chow or semi-synthetic diets (SSD), and administered either proanthocyanidins (PAC) or water (control). Adult and larval *T. muris* counts (**B-C**), total cellularity of the mesenteric lymph nodes (**D**), proportions of T-bet^+^ (Th1) and GATA3^+^ (Th2) cells within the MLN TCRβ^+^ CD4^+^ population, and the Th1/Th2 ratio (**E-G**), serum IgG2a specific for *T. muris* antigen (**H**), and IL-6 secretion from MLN cells stimulated with *T. muris* antigen (**I**) at day 35 following the start of *T. muris* infection. n= 6 per treatment group from a single experiment. \**p* < 0.05, \*\**p* < 0.01, \*\*\**p*<0.005 by two-way ANOVA with Tukey post-hoc testing. Shown are means ± S.E.M.

### Proanthocyanidins and helminths interact to change gut microbiota composition

We next assessed if PAC intake and/or helminth infection influenced GM composition. In the *H. polygyrus* infection model, PAC intake significantly increased α-diversity in the caecal microbiota based on the number of observed zOTUs, but not on Shannon diversity index, whilst *H. polygyrus* infection had no effect on these measures (**Figure 5A**). Pairwise PERMANOVA analysis assessing cecal microbiota compositional differences (Bray-Curtis dissimilarity based) revealed a clear interaction between *H. polygyrus* infection and PAC intake, whereby the effect of PAC was modulated by concurrent infection. In uninfected mice, the caecal microbiome composition was significantly different in mice fed PAC relative to the control diet (adjusted *p* < 0.05; **Figure 5B**). In contrast, PAC had no effect in *H. polygyrus* -infected mice (adjusted *p* > 0.05), indicating that infection abrogated the effects of PAC intake on the caecal microbiome composition. In contrast, regardless of PAC intake, *H. polygyrus* infection resulted in significant changes in the caecal microbiome composition relative to uninfected mice, (adjusted *p* < 0.05; **Figure 5B**). Pairwise DESeq2 analysis identified 10 and 12 species that were impacted by *H. polygyrus* infection in control- and PAC-dosed mice, respectively (Supplementary Figure 5). Similar to previous studies^39, 40^, the major change induced by *H. polygyrus* was an increase in lactobacilli, predominately *Lactobacillus johnsonii,* and *Bifidobacterium animalis* subsp. *lactis (p* <0.05 for main effect of infection by two-way ANOVA; **Figure 5C**). In uninfected mice, PAC intake resulted in three differentially abundant taxa by DESeq2, these being a reduction in the relative abundance of *Ligilactobacillus animalis* and increases within unclassified members of the *Gordonibacter* and *Corynebacterium* members (Supplementary Figure 5). Moreover, PAC intake tended to increase (adjusted *p*=0.06 by DESeq2) the abundance of *Turicibacter sanguinis,* a bacterium associated with altered fat digestion and reduced triglyceride levels^41^, consistent with reported effects of PAC-rich diets on lipid metabolism^42^. However, these effects of PAC were strongly influenced by concurrent *H. polygyrus* infection, with relative *L. animalis* abundance not affected by PAC in infected mice (**Figure 5C**). Moreover, infection strongly suppressed *T. sanguinis* abundance and completely abrogated the PAC-induced increase in *T. sanguinis*. Indeed, within *H. polygyrus*-infected mice, *T. sanguinis* relative abundance was significantly lower in PAC-dosed mice (*p* < 0.05 for interaction between diet and infection by two-way ANOVA; **Figure 5C**). Collectively, these data indicate that PAC-induced changes in GM composition were attenuated by concurrent *H. polygyrus* infection.

**Figure 5.**
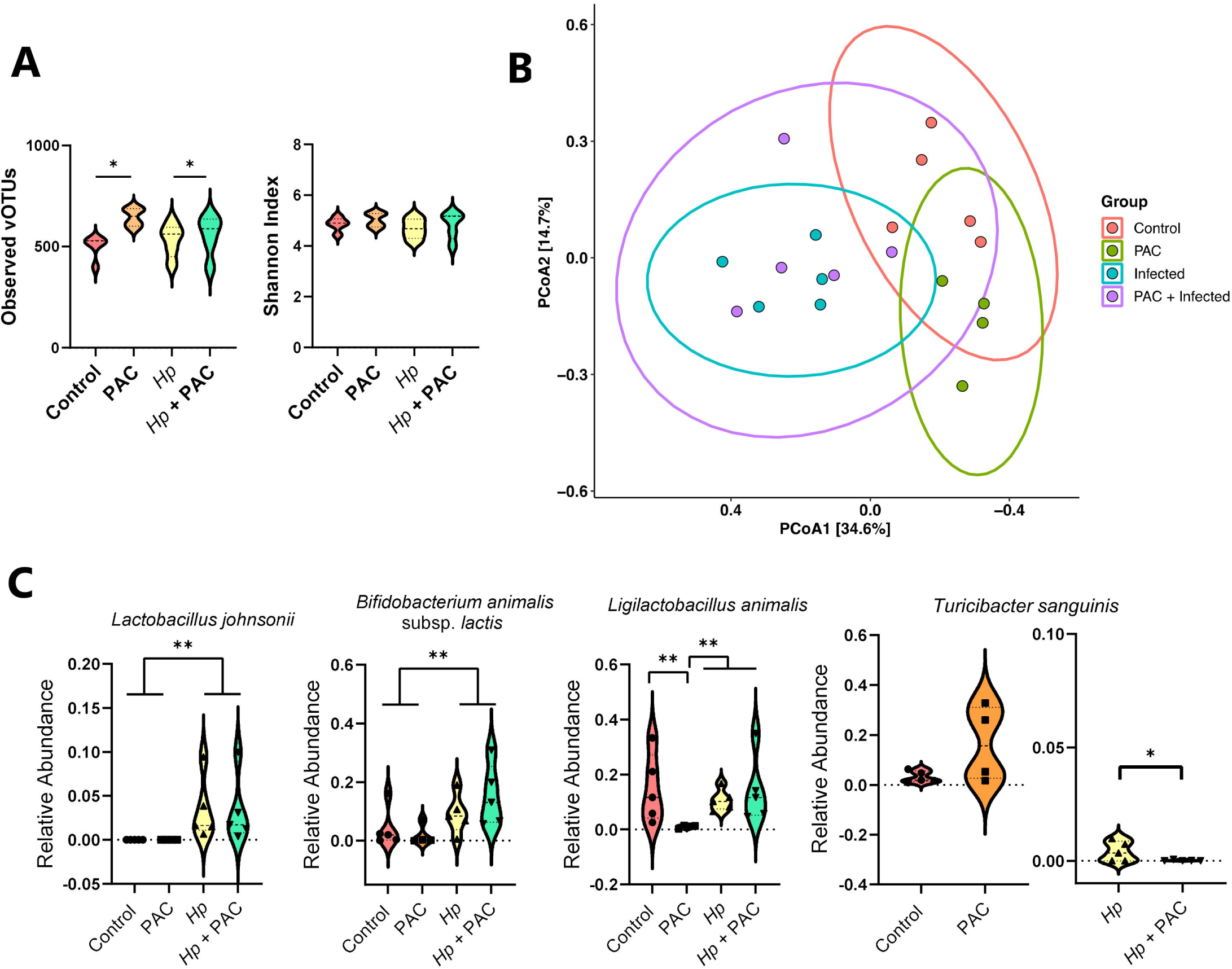
Impact of *Heligmosomoides polygyrus* (*Hp*) infection and proanthocyanidins (PAC) on the host caecal microbiota. **A)** Violin plots illustrating the number of observed zOTUs and Shannon Diversity Index for each treatment group. * *p*< 0.05 by two-way ANOVA followed byTukey post-hoc testing. **B)** Principal coordinates analysis based on Bray-Curtis dissimilarity metrics for control and *Hp*-infected mice dosed with either PAC or water (control). Each data point on the PCoA plots indicate a sample, with ellipses showing 95% confidence interval. The percentage in brackets is the percentage of variation explained by each PCoA axis. Pairwise comparisons between treatment groups were performed using permutation MANOVAs on a distance matrix with p-value correction using the Holm method. *p*<0.05 was considered as significant. **C)** Relative abundance bar plot for *Lactobacillus johnsonii*, *Turicibacter sanguinis*, *Ligilactobacillus animalis,* and *Bifidobacterium animalis* subsp. *lactis* in uninfected and *Hp*-infected mice dosed with either PAC or water (control) (n= 4-5 per treatment group). The significance of difference was assessed by two-way ANOVA and Tukey post-hoc testing (* *p*<0.05; \*\**p* < 0.01).

In faecal samples of *T. muris* -infected mice (35 days post-infection), infection had the strongest effect on GM composition. Infected mice had significantly lower α-diversity (both in terms of observed zOTUs and Shannon index metrics), whilst PAC had no effect (**Figure 6A**). The faecal microbiome compositions of infected mice tended to diverge from uninfected mice, regardless of PAC intake (adjusted *p* = 0.08 for effect of infection in both control-fed and PAC-fed mice by pairwise PERMANOVA on Bray-Curtis dissimilarity metrics; **Figure 6B)**. DESeq2 analysis indicated that infection changed the abundance of 15 and 17 species in control and PAC-dosed mice, respectively (Supplementary Figure 6). Notably, when comparing the overall effect of infection (irrespective of PAC intake) there was a significant increase in *Li. animalis* abundance, consistent with previous studies showing an expansion in lactobacilli in mice with a chronic *T. muris* infection^43^. The effect of PAC on faecal GM composition was not as pronounced as observed in the *H. polygrus* experiments, which may relate to the differences in samples analyzed (caecum vs. faeces) or the length of study (28 vs 49 days). Despite this, we once more noted a non-significant trend for a relative enrichment of *T. sanguinis* in uninfected, PAC-dosed mice (**Figure 6C**) despite a significant reduction in PAC-dosed mice during *T. muris* infection, again suggesting that infection abrogated the effects of PAC as was observed also for *H. polygyrus* infection (*p* = 0.05 for interaction between diet and infection by two-way ANOVA; **Figure 6C**). Interestingly, we also found that the combination of PAC and *T. muris* infection led to the expansion of the pathobiont *Escherichia fergusonii* which was absent in all other treatment groups (*p* <0.01 for interaction between diet and infection by two-way ANOVA; **Figure 6C)**. Thus, consistent with the immunological data, administration of PAC to *T. muris* -infected mice tended to result in an increase in parameters associated with inflammation. Collectively, these data indicate that significant interactions exist between gut parasites and dietary PAC on GM composition.

**Figure 6.**
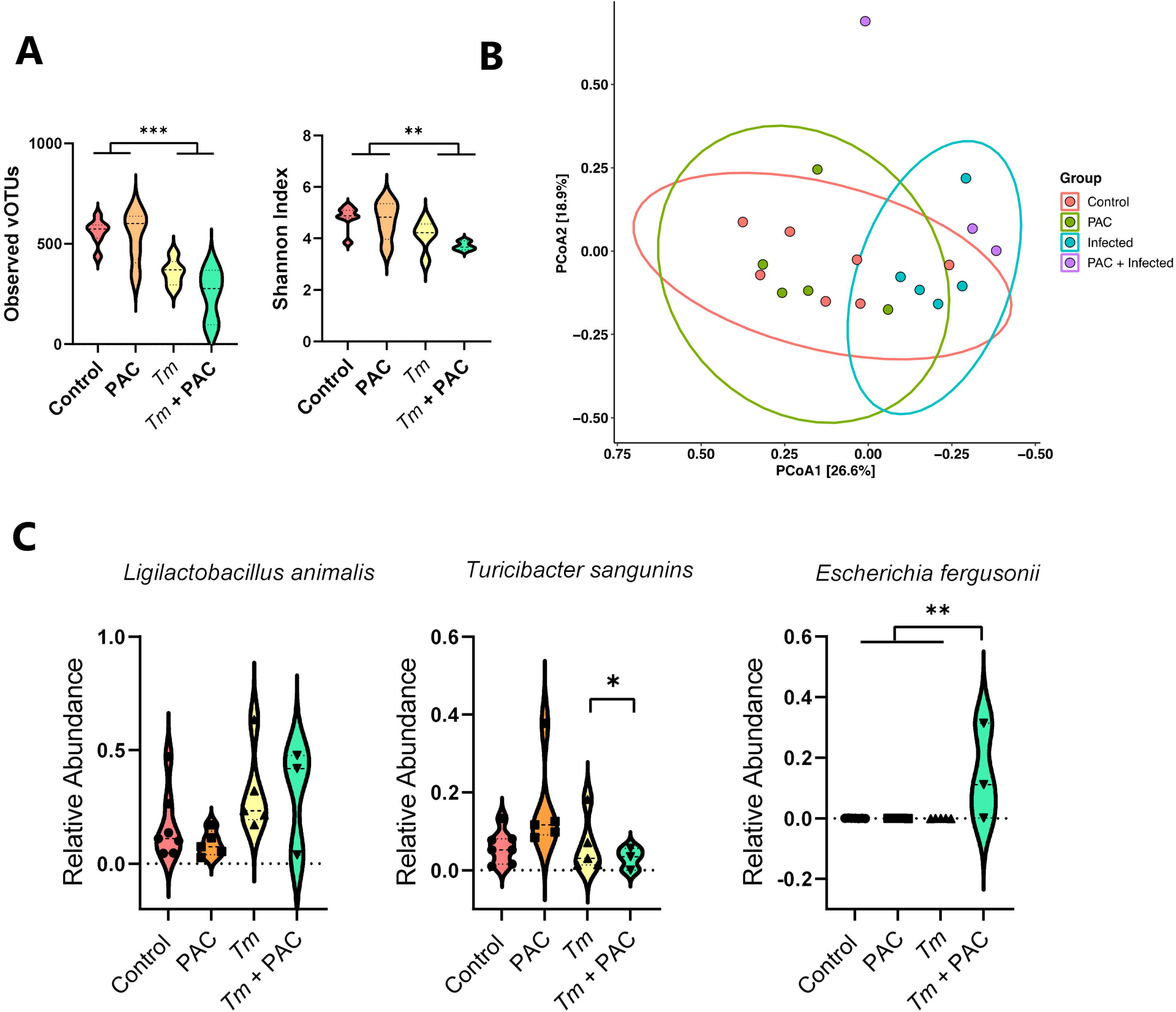
Impact of *Trichuris muris* (*Tm*) infection and proanthocyanidins (PAC) on the host faecal microbiota. **A)** Violin plots illustrating the number of observed zOTUs and Shannon Diversity Index for each treatment group. *** *p*< 0.001, ** *p*<0.01 by two-way ANOVA followed by Tukey post-hoc testing. **B)** Principal coordinates analysis based on Bray-Curtis dissimilarity metrics for control and *Tm*-infected mice dosed with either PAC or water (control). Each data point on the PCoA plots indicate a sample, with ellipses showing 95% confidence interval. The percentage in brackets is the percentage of variation explained by each PCoA axis. No ellipse was drawn for the *Tm*-infected mice dosed with PAC group as n=3. Pairwise comparisons between treatment groups were performed using permutation MANOVAs on a distance matrix with p-value correction using the Holm method. *p*<0.05 was considered as significant. **C)** Relative abundance bar plots for *Ligilactobacillus animalis, Turicibacter sanguinis*, and *Escherichia fergusonii* in uninfected and *T. muris*-infected mice dosed with either PAC or water (control) (n= 3-7 per treatment group). **p < 0.01, \**p<*0.05 by Tukey post-hoc testing following two-way ANOVA.

### *Trichuris muris* infection changes the profile of microbial metabolites derived from proanthocyanidins

Given that helminth infection appeared to influence how the GM responded to PAC supplementation, we next assessed how *T. muris* and PAC may interact to influence the production of GM-derived metabolites. To this end, faecal samples of uninfected and *T. muris-*infected mice administered PAC were analyzed for short chain fatty acid (SCFA) concentrations. We observed significant interactions (*p* < 0.05) between PAC intake and infection for acetic acid, propionic acid, and total SCFA. These interactions reflected a generally higher level of SCFA in either uninfected PAC-dosed mice or *T. muris* -infected control mice, but a lower level in the combinatorial group of PAC-dosed *T. muris* -infected mice (**Figure 7A)**. Thus, whilst both treatments in isolation promoted SCFA production, instead of an additive effect we detected an antagonistic trend. Butyric acid followed the same trend but was not significant (**Figure 7A)**.

**Figure 7.**
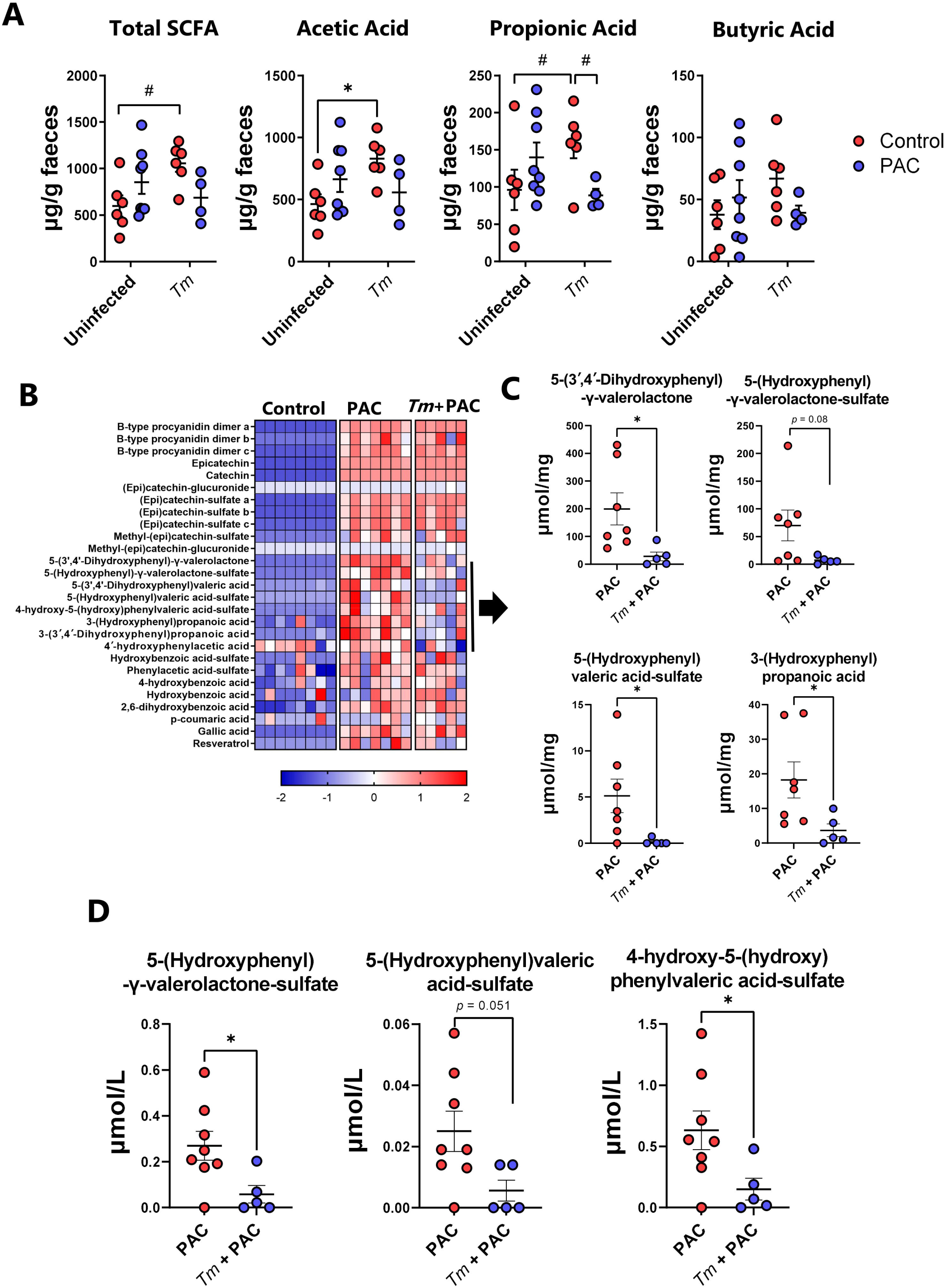
Short chain fatty acids and identification of proanthocyanidin metabolites in *Trichuris muris* infected mice. **A)** Effect of proanthocyanidins (PAC) and *Trichuris muris* (Tm) infection on the concentrations of total short chain fatty acids (SCFA), acetic acid, propionic acid, and butyric acid in faecal samples. A significant (*p* < 0.05) interaction between diet and infection was noted for acetic acid, propionic acid and total SCFA by two-way ANOVA. **B)** Identification of PAC metabolites in the caecum of PAC-dosed uninfected and *T. muris*-infected mice relative to uninfected mice dosed with water. **C)** Caecal and **D)** serum concentrations of PAC-derived metabolites in PAC-dosed uninfected and *T. muris* infected mice. n=5-8 per group, pooled from two independent experiments. Shown are means ± S.E.M. or median values. * *p*<0.05 by unpaired t-test.

Several studies have shown that absorption of PAC in the small intestinal is minimal, while they are readily metabolized by the residing GM in the large intestine. Accordingly, PAC-derived phenolic metabolites such as phenyl-γ-valerolactones have been proposed to play a role in their putative health benefits, such as improved vascular function^44, 45^. Therefore, we used targeted metabolomics to examine whether phenolic metabolites produced from the breakdown of PAC were altered during *T. muris* infection, by quantifying the abundance of a panel of PAC-derived metabolites in the caecum of uninfected and *T. muris*-infected mice dosed with PAC. As expected, PAC supplementation resulted in substantial increases of these metabolites in the caecum, compared to control-fed mice. Most of the metabolites were similar in both groups of mice that received PAC supplementation, regardless of infection (**Figure 7B)**. However, we identified a cluster of related metabolites where abundance was reduced in infected mice. These were mostly phenyl-γ-valerolactones and phenylvaleric acid derivatives including 5-(31,41-dihydroxyphenyl)-γ-valerolactone, 5-(hydroxyphenyl)-γ-valerolactone-sulfate, 5-(hydroxyphenyl)valeric acid-sulfate, and 3-(hydroxyphenyl)propionic acid (**Figure 7C)**. Moreover the serum concentration of 5- (hydroxyphenyl)-γ-valerolactone-sulfate and 4-hydroxy-5-(hydroxyphenyl)valeric acid-sulfate were also significantly reduced in infected mice dosed with PAC (**Figure 7D).** 3-(Hydroxyphenyl)propionic acid and 5-(31,41-dihydroxyphenyl)-γ-valerolactone were not detected in the serum in any of the treatment groups (data not shown). Thus, both *T. muris* and PAC supplementation in isolation appeared to have positive effects on the production of microbial-derived metabolites in the gut, but concurrent infection and PAC intake attenuated both SCFA levels and the production and absorption of phenyl-γ-valerolactones, suggestive of an antagonistic interaction that may have a negative effect on tissue homeostasis and intestinal health.

## Discussion

Whilst the immunomodulatory effects of parasites have been extensively demonstrated in previous studies, our understanding of the interactions between infection and bioactive dietary components remains limited^46, 47^. Similarly, despite numerous investigations into the impact of helminth infection on the GM, diet-parasite interactions and their effects on gut health are largely unexplored^39, 48, 49^. Several studies have demonstrated strong anti-inflammatory properties of PAC, which have also been shown to be advantageous for gut health by supporting mucosal barrier function^6, 50^. Moreover, PAC may alter gut microbiota by enhancing the growth of gut commensals, such as *Bacteroides* species^51–53^. Thus, by altering the bacterial flora of the intestinal tract, PAC may indirectly stimulate gut-associated myeloid tissues, and thereby modulate T- or B-cell mediated immune responses^54, 55^.

Based on these known anti-inflammatory and immunomodulatory properties, we hypothesized that PAC may reduce helminth-induced inflammation by either enhancing the diversity and composition of the GM, or by directly stimulating mucosal immune cells. Both infection models clearly showed a higher number of MLN cells in infected mice dosed with PAC, which suggests a strong effect of PAC on immune reactivity and lymphocyte proliferation. However, we found inconsistent effects of PAC on the immune polarization during *H. polygyrus* infection, but a more pronounced type-1 polarized immune response in PAC-treated *T. muris-*infected mice. The differences between the models may relate to the duration of the experiments, the different tissue location of the two parasites, or perhaps intrinsic differences in the host-parasite relationships. However, it was notable that, in the absence of infection, PAC had a much stronger effect on the caecal transcriptional response than the small intestinal response, which may suggest that the large intestine is more responsive to plant-derived phytochemicals such as polyphenols.

As PAC are known to be metabolized in the large intestines, we sought to identify which PAC metabolites were present in the caecum. As expected, several PAC metabolites were identified in PAC-treated mice. However, significantly fewer metabolites were found in *T. muris* infected mice dosed with PAC, including valerolactones. This may be closely related to the changes in the GM composition induced by helminth infection, which may alter PAC metabolism efficiency. Another suggestion could be that PAC metabolites are more easily absorbed in the gut, due to the increased permeability and disruption of the gut mucosal barrier caused by the helminth infections. However, the serum concentration of valerolactones was also significantly lower in infected mice, suggesting that impaired production of these metabolites by the GM seems most plausible.

Semi-synthetic diets resemble Western-style diets that lack crude fibrous plant components such as pectin, lignin, and (poly)phenols. The consumption of such diets also often contains elevated levels of fat and simple sugars and is associated with the development of diseases such as obesity and colitis. It has been shown that incorporation of plant-derived fibers or phytochemicals such as PAC into refined, high fat diets can be protective against disease in mice^33, 56^. However, the inclusion of high levels of fermentable fibers in refined diets has been shown to also have some detrimental effects, such as worsening of acute inflammation deriving from colitis induced by chemicals (DSS)^37^ or infectious agents (*C. rodentium*)^57^. Notably, we have recently shown that high levels of dietary inulin (a prebiotic oligosaccharide with known anti-inflammatory effects) also impaired immunity to *T. muris*, with higher worm burdens and a Th1 driven immune response, indicating an important context-dependent immunological effect of fermentable fiber^58^. Together with our current results, this suggests that caution must be exercised when fortifying refined diets with high levels of phytochemicals during active enteric inflammation or infection. Indeed, mouse chow, which contains a high level of crude, unrefined plant fiber, also substantially increased *T. muris* burdens, suggesting a continuum whereby increasing the concentration of prebiotic substrates progressively increases susceptibility to parasites. Thus, whilst the well-known health benefits of PAC or other plant fibers are clear, their immunomodulatory effects may differ according to context.

Therefore, the lack of a complex GM, which could also be expected to be found in mice fed SSD with little plant material or fermentable fibers, seems to promote resistance to infection whilst the addition of PAC, inulin, or the feeding of unrefined chow progressively down-regulates type-2 immunity to infection. Thus, the production of hitherto undefined GM-derived metabolites likely shapes the propensity of immune cells to diverge away from type-2 functionality to either type-1 or T-regulatory responses, which may, in some cases, predispose to enteric infections. Thus, while our results and others clearly indicate that even if PAC and other phytonutrients have demonstrated beneficial effects on gut health in the absences of gut pathogens, further studies should aim at unravelling the immunological mechanisms underlying this complex relationship between diet and immune function during infection with gastrointestinal parasites.

## Material and methods

### Proanthocyanidins

PAC derived from grape (*Vitus vinifera)* consisting of a mixture of oligomers and polymers were used for all experiments. For *H. polygyrus* experiments, PAC were derived from grape pomace and obtained by series of extraction and sephadex separation LH-20 gel chromatography as described previously^59^. The PAC had a mean degree of polymerization of 7.5. PAC used for the *T. muris* study were a commercial preparation derived from grape seed (Bulk Powders, Denmark), with a mean degree of polymerization of 4.2. Purity of both preparations was >95% as determined by LC-DAD-MS and LC-DAD-MS/MS analyses^60^.

### Parasites

*H. polygyrus* and *T. muris* were propagated, and excretory/secretory (E/S) antigens produced, as described previously^58, 61^.

### Mouse experiments

6-week-old C57/BL6 female mice were used in all experiments, and all mice were fed a purified control diet (E15051-04, Ssniff, Germany, composition in Supplementary Table 1) throughout the entire study period. All mice were given 1 week acclimatization, were monitored daily and weighed once a week. The mice were subjected to a 12 h light/dark cycle (6:00 a.m. to 6:00 p.m.) with *ad libitum* access to water and food, and randomly assigned to treatment groups. For *H. polygyrus* experiments, mice were orally gavaged on alternate days for 4 weeks with either PAC dissolved in sterile water (200 mg/kg BW, PAC, or *H. polygyrus +* PAC groups) or water (control and *H. polygyrus* groups). Mice were infected with *H. polygyrus* (200 third-stage larvae/mouse in 200 µl water) on day 14 and were humanely euthanized on day 28 (i.e., 14 days p.i.). For *T. muris* experiments, all mice were orally gavaged on alternate days for 7 weeks with either PAC (300 mg/kg body weight, PAC and *T. muris* + PAC groups) or water only (control and *T. muris* groups). After 2 weeks of PAC treatment, the appropriate treatment groups were infected with 20 eggs of *T. muris* on day 0, 14 and 28 to establish naturally occurring chronic infection^62^. At the end of the experimental period (day 35 post first infection), all mice were humanely euthanized by cervical dislocation.

### Sample collection

Fecal samples and blood were collected at necropsy. Separated serum was stored at -20 °C until use. After sacrifice, the MLN were dissected and stored on ice in 10 % fetal calf serum (FCS)- supplemented RPMI 1640 media (complete media) until further processing for flow cytometry. Caecal contents were harvested and cooled immediately on ice and stored at – 80 °C until used for extraction. Tissue samples from the duodenum of *H. polygyrus* infected mice, and the caecal tip of *T. muris* infected mice, were collected and stored in RNA later. Full-thickness duodenum samples were collected from *H. polygyrus* -infected mice for histology. Histology samples were longitudinally opened, gently rinsed with PBS, and stored in 4 % paraformaldehyde until further processing. Worm burdens were assessed by manual enumeration under a stereomicroscope.

### Histology

Samples stored in 4% paraformaldehyde were embedded in paraffin blocks, sectioned, and mounted on glass slides prior to Periodic-acid Schiff (PAS) or Hematoxylin and eosin (H&E) staining. Furthermore, Mcpt1-positive mast cells were stained in additional paraffin-embedded sections, which were de-waxed, and Ag retrieval was performed with citrate buffer. Tissue sections were incubated with rat anti-mouse primary monoclonal mast cell protease-1 (MCPT-1) Ab (1:100, clone RF6.1; Thermo Fisher Scientific), followed by secondary staining with biotinylated rabbit anti-rat IgG (Abcam). Mcpt1^+^ cells were manually enumerated by a microscopist blinded to the treatment groups.

### Isolation of mesenteric lymph node cells

MLN were carefully dissected and trimmed of fat. Single cell suspensions were prepared by passing through a 70 μM cell strainer. Afterwards, the cell suspension was centrifuged at 450g for 5 minutes at room temperature. Cell counts and viability were assessed manually using a haematocytometer and trypan blue staining.

### Flow cytometry, ex vivo cell stimulation and cytokine analysis

MLN cell suspensions were incubated with Fc-block (anti-CD16/CD32, BD Biosciences, cat no. 553142) followed by staining with antibodies against surface and intranuclear markers. Cells were kept at 4°C throughout the staining procedure. Cell surface markers were stained for 20 min using an antibody cocktail containing; FITC-conjugated hamster anti-mouse TCRβ (clone H57-597; BD Biosciences, cat no. 553171), and PerCP-Cy5.5-conjugated rat anti-CD4 (RM4-5; BD Biosciences cat no.550954). For intranuclear (GATA-3 and T-bet) staining, FoxP3/Transcription Factor staining buffer (eBiosciences, 00-5523-00) was used according to manufacturer’s instructions. Fixed/ permeabilized cells were incubated for 30 min on ice with: Alexa Fluor® 647-conjugated mouse anti-mouse T-bet (4B10; BD Biosciences cat no. 561264), PE-conjugated rat anti-mouse GATA3 (TWAJ; Thermo Fisher Scientific cat no. 12-9966-42), FITC-conjugated rat anti-mouse FoxP3 (FJK-16s; Thermo Fisher Scientific cat no. 11-5773-82). Cells were analyzed on an BD Accuri C6 flow cytometer (BD Biosciences). All data were acquired and analyzed using Accuri CFlow Plus software (Accuri^®^ Cytometers Inc.). Gating strategy is shown in Supplementary Figure 7.

For cytokine analysis, MLN cells were plated in triplicate into 96-well cell culture plates at the density of 5.0 x 10^6^ cells/mL in complete media and stimulated with *T. muris* E/S antigens (50 µg/mL) or PBS. Cells were incubated at 37 °C/5 % CO_2_ and cell-free supernatants were harvested after 24 hours and stored at -20°C for subsequent analyses. Secreted cytokines were measured using a BD Th1/Th2/ Th17 cytometric bead array (CBA) kit (BD Biosciences cat no. 560485) according to manufacturer’s instructions. Samples were processed on a BD Accuri C6 flow cytometer (BD Biosciences), with data acquired using Accuri CFlow Plus software (Accuri® Cytometers Inc., MI, USA).

### Enzyme-linked immunosorbent assay

*T. muris* E/S-specific antibodies were measured from diluted (1:50) serum using a previously described ELISA protocol^58^. The antibodies employed for ELISA were biotin-conjugated rat anti-mouse IgG2a (clone R19-5, BD Biosciences, Denmark cat no. 550332) and anti-mouse IgG conjugated to horseradish peroxidase (HRP; Bio-Rad, Germany cat no. 1706516). Absorbance was measured at an optical density of 450 nm with a Multiskan FC plate reader (Waltham, MA, USA). *H. polygyrus* E/S-specific IgG1 was detected using goat anti-mouse IgG1-HRP conjugate (Invitrogen, cat no. A10551).

### RNA extraction, RNA sequencing, and qPCR

Tissue was mechanically homogenized in QIAzol lysis buffer using a gentleMACS™ dissociator (Miltenyi Biotec, Germany) and total RNA was isolated using miRNeasy Mini Kits (Qiagen, CA, USA) as per manufacturer’s instructions. Total RNA concentrations were determined using a NanoDrop ND-1000 spectrophotometer (NanoDrop Technologies, DE, USA). Paired-end (100 bp) RNA-sequencing was carried out using the DNBSEQ sequencing platform (BGI, Copenhagen, Denmark). Clean reads were mapped to the mouse genome (mm10) using Bowtie2 (v2.2.5). Differentially expressed genes were detected using DEseq2^63^. cDNA was synthesized from 500 ng of RNA using Quantitect Reverse Transcriptase kits (Qiagen) and qPCR was performed using perfeCTa SYBR green fastmix (Quanta Bioscience) using the following program: 95°C for 2 minutes followed by 40 cycles of 15 seconds at 95°C and 20 seconds at 60°C. Primer sequences are listed in Supplementary Table 2.

### Short chain fatty acids and proanthocyanidin metabolites analysis

SCFA were measured by GC-FID using the same methodology as described by Choi *et al*.^64^, whilst PAC metabolites were analyzed using UHPLC-QToF using previous described methods. For PAC metabolites analysis, sample preparation of plasma was done according to the protocol of Dudonné *et al.* ^65^. For the caecum, samples were freeze-dried before being extracted with 25 µL per mg of dry matter of methanol:water (50:50 v/v) spiked with 0.5 ppm of 4-hydroxybenzoic acid-d_4_. Samples were homogenenized for 2 min with a Bead Rupter12 (Omni International, Kennesaw, GA, USA), vortexed during 5 min, sonicated for 30 min at room temperature and re-vortexed during 5 min. Before injection, samples were centrifuged at 18 000 g during 30 min at 4 °C before being filtered through a 0.22 µm Nylon filter. PAC metabolites analysis was carried out with an Acquity I-Class UHPLC coupled with a Synapt G2-Si (Waters, Milford, MA) using the UHPLC-QToF method from Lessard-Lord *et al*.^66^. PAC and (epi)catechin derivatives were quantified as epicatechin equivalent and PAC metabolites as 5-(31,41-dihydroxyphenyl)-γ-valerolactone equivalent, while gallic acid was quantified with its own standard.

### Microbiota analysis

#### DNA extraction and amplification of the 16S rRNA gene V3 region

DNA was extracted from 100mg caecal content (*H. polygyrus* study) and faecal samples (*T. muris* study) using the Bead-Beat Micro AX Gravity kit (A&A Biotechnology, Poland) as per manufacturer’s instructions with the addition of mutanolysin and lysozyme to enhance the bacteria lysis yield. The concentration of DNA was assessed utilizing the Qubit® dsDNA HS Assay Kit (Life Technologies, CA, USA) and measurement was taken using the Varioskan Flash Multimode Reader (Thermo Fisher Scientific, MA, USA). Library was prepared through a two-step PCR process. In the initial PCR step, the 16S rRNA gene V3 region was amplified using the Nextera Index Kit (Illumina, CA, USA) compatible forward primer nxt388_F:(5’-TCGTCGGCAG CGTCAGATGT GTATAAGAGA CAGACWCCTA CGGGWGGCAGCAG-3’) and reverse primer nxt518_R:(5’-GTCTCGTGGG CTCGGAGATG TGTATAAGAG ACAGATTACC GCGGCTGCTGG-3’). This step utilized a final reaction volume of 25 μl/sample, incorporating 5 μl of 51×1PCRBIO HiFi buffer (PCR Biosystems^©^, UK), 0.25 μl of PCRBIO HiFi Polymerase (PCR Biosystems^©^, UK), 0.5 μl of primer mix (10μM, nxt_388_F and nxt_518_R), 5 μl of genomic DNA (5ng/μl), and was adjusted to 25 μl with nuclease-free water. Reaction conditions included an initial denaturation at 95°C for 2 min, followed by 33 cycles of: 95°C for 15 seconds, 55°C for 15 seconds and 72°C for 20 seconds, and a final extension step at 72°C for 4 min. A second PCR step was conducted, involving 5 μl of 51×1PCRBIO HiFi buffer (PCR Biosystems^©^, UK), 0.25 μl of PCRBIO HiFi Polymerase (PCR Biosystems^©^, UK), 2 μl of each P5 and P7 primer (Nextera Index Kit, Illumina, CA, USA), 2 μl of initial PCR product, and additional nuclease-free water to make up a total volume of 25 μl. Reaction parameters of PCR2 consisted of an initial denaturation at 95°C for 1 min, followed by 13 cycles of: 95°C for 15 seconds, 55°C for 15 seconds and 72°C for 15 seconds, and a final extension step at 72°C for 5 min. The second PCR products were cleaned up using SpeedBeads™ magnetic carboxylate (obtained from Sigma Aldrich). Cleaned PCR2 products were quantified using Qubit® dsDNA HS Assay Kit (Life Technologies, CA, USA) and then mixed equimolarly. High-throughput sequencing of 16S rRNA gene amplicon (V3 region) was done, using a NextSeq550 platform (Illumina, San Diego, CA, USA) with the Mid Output Kit v2 (300 cycles), to determine the bacterial composition of caecal and faecal contents.

#### Data processing and analysis

The merging and trimming of raw dataset containing pair-ended reads with corresponding quality scores were carried out using *fastq_mergepairs* and *fastq_filter* scripts implemented in the VSEARCH pipeline^67^ utilizing the specified settings: -fastq_minovlen 100, -fastq_maxee 2.0 - fastq_truncqual 4 -fastq_minlen 150. Removal of chimeric reads from dataset and constructing zero-radius operational taxonomical units (zOTUs) were performed by using UNOISE3 in VSEARCH pipeline. The Greengenes (13.8)^68^ 16S rRNA gene collection was used as a reference database and taxonomical assignments were obtained by using SINTAX^69^ for the 16S rRNA gene database. zOTUs belonging to bacteria assigned at phylum level was included in analysis. Unassigned taxonomy names at any levels lower than phylum were adjusted to be termed as “Unclassified *lowest taxonomy name*”. zOTUs mapped to *Cyanobacteria*/Chloroplast were excluded prior to the analysis. The taxonomy names of zOTUs previously belonging to the genus *Lactobacillus* were updated manually following the announcement of Zheng et al.^70^.

The analysis and visualization of data from the two studies conducted for each infection model were performed independently in RStudio using R version 4.2.1 and R packages phyloseq^71^, DESeq2^63^, tidyverse^72^, reshape2^73^, pheatmap^74^, and RColorbrewer^75^. Alpha and beta diversity analyses were performed on rarefied data (7000 reads/sample). Alpha diversity was assessed by observed zOTUs and Shannon Index metrics, while beta diversity was evaluated through principal coordinates analysis (PCoA) with Bray–Curtis dissimilarity metrics. A DESeq2 analysis was conducted to determine species level differentially abundant bacteria between treatments and DESeq2-based heatmap was plotted to show significantly differentially abundant bacteria with an adjusted *p* value below a threshold of 0.05. In addition, within each study, bar plots were drawn to show how the relative abundance of selected bacteria identified by DESeq2 changed among some of the treatment groups.

For beta diversity, pairwise comparisons between treatment groups within each study were conducted by using permutation MANOVAs on a distance matrix with Holm *p* value adjustment method using the *pairwise.perm.manova* function available in R RVAideMemoire package^76^. In DESeq2 analysis, *p* values attained by the Wald test were corrected by using the Benjamini and Hochberg method by default, and adjusted *p* value threshold of below 0.05 was considered as significant.

### Statistical analysis

Statistical analyses were performed in Prism 8.0.2 (GraphPad Software). Assumptions of normality were checked through Shapiro-Wilk tests, or inspection of histogram plots and Kolmogorov-Smirnov tests of ANOVA residuals. Parametric data were analyzed using unpaired t-tests, or ANOVA followed by Tukey post-hoc testing, and presented as means ± S.E.M. Non-parametric data were analyzed using Mann-Whitney tests or Kruskal-Wallis and Dunn’s post hoc tests, and results presented as median values. Details of each experiment are given in the appropriate figure legends.

## Acknowledgements

The authors would like to thank Mette Marie Arnt Schjelde and Penille Jensen for excellent laboratory assistance and support for the conduction of animal studies. We are grateful to Rick Maizels (University of Glasgow) and Sebastian Rausch (Freie University, Berlin), for provision and advice on *H. polygyrus* infections. This work was funded by Independent Research Fund Denmark (Grant # 702649B).

## Ethical statement

Mice experiments were approved by the Danish Animal Experiment Inspectorate (Ref. No. 2015-15-0201-00760) and performed in accordance with EU Directive 2010/63/EU for animal experiments.

## Data availability

RNA sequence data from caecum and duodenum are deposited at the NCBI Gene Expression Omnibus (GEO: Accession numbers GSE174756 and GSE176182). The raw 16S rRNA sequencing data can be accessed at Sequence Read Archive (https://www.ncbi.nlm.nih.gov/sra) using the accession number PRJNA1044063.

