## Supplementary Table and Figures for "Gut microbial-mediated polyphenol metabolism is restrained by parasitic whipworm infection and associated with altered immune function in mice"

**Contents:**

Tables S1-S2

Figures S1-S7

Files S1-S6

**Supplementary Table 1 –** Composition of purified basal diet (semi-synthetic diet)

| **Feed component (g/100g)** | |
|---|---|
| Casein | 20.0 |
| Corn starch | 37.6 |
| Maltodextrin | 15.0 |
| Sucrose | 10.0 |
| Cellulose powder | 5.0 |
| L-Cysteine | 0.2 |
| Vitamin premix | 1.0 |
| Mineral & trace element mix | 6.0 |
| Choline chloride | 0.2 |
| Soybean oil | 5.0 |
| **Crude Nutrients (%)** | |
| Crude protein | 17.9 |
| Crude fat | 5.1 |
| Crude fiber | 5.0 |
| Crude ash | 5.4 |
| Starch | 36.2 |
| Sugar | 11.0 |
| **Energy (MJ [or kcal] ME/kg)** | 15.4 |

**Supplementary Table 2 –** Primers used for qPCR

| Gene | Forward Primer (5’ – 3’) | Reverse Primer (5’ -3’) |
|---|---|---|
| *Mcpt1* | CTCTGCATATGTGCCCTGGATTA | GGGGGCAGACTGGGGATAGT |
| *Ifit3b* | TTCCCAGCAGCACAGAAACA | GTTGCACACCCTGTCTTCCA |
| *Irgm1* | CCATGCGTATGCTGCTGTTT | GATCTGCGGAGGGAAGATGG |
| *Gapdh* | TATGTCGTGGAGTCTACTGGT | GAGTTGTCATATTTCTCGTGG |

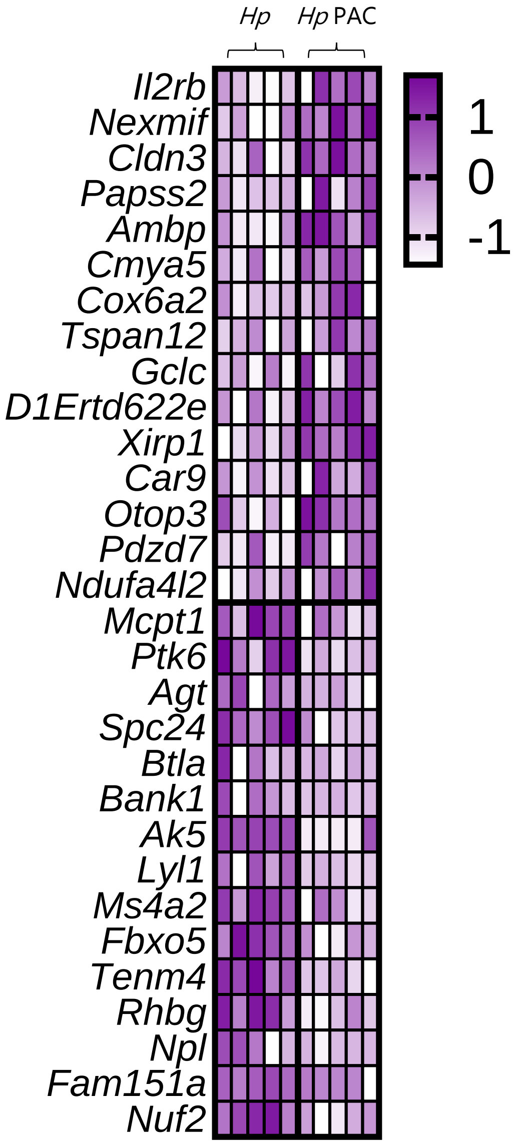


| Gene | padj | p value |
|---|---|---|
| Il2rb | 0.999726 | 9.92E-05 |
| Nexmif | 0.999726 | 0.001475 |
| Cldn3 | 0.999726 | 0.001716 |
| Papss2 | 0.999726 | 0.001778 |
| Ambp | 0.999726 | 0.00194 |
| Cmya5 | 0.999726 | 0.002187 |
| Cox6a2 | 0.999726 | 0.002571 |
| Tspan12 | 0.999726 | 0.003658 |
| Gclc | 0.999726 | 0.004774 |
| D1Ertd622e | 0.999726 | 0.004904 |
| Xirp1 | 0.999726 | 0.005543 |
| Car9 | 0.999726 | 0.006459 |
| Otop3 | 0.999726 | 0.006619 |
| Pdzd7 | 0.999726 | 0.006817 |
| Ndufa4l2 | 0.999726 | 0.006977 |
| Mcpt1 | 0.999726 | 0.003658 |
| Ptk6 | 0.999726 | 0.003484 |
| Agt | 0.999726 | 0.003416 |
| Spc24 | 0.999726 | 0.003389 |
| Btla | 0.999726 | 0.003273 |
| Bank1 | 0.999726 | 0.002559 |
| Ak5 | 0.999726 | 0.002506 |
| Lyl1 | 0.999726 | 0.002426 |
| Ms4a2 | 0.999726 | 0.001998 |
| Fbxo5 | 0.999726 | 0.001309 |
| Tenm4 | 0.999726 | 0.001004 |
| Rhbg | 0.999726 | 0.000631 |
| Npl | 0.999726 | 0.00062 |
| Fam151a | 0.999726 | 0.000533 |
| Nuf2 | 0.999726 | 0.000216 |

**Supplementary Figure 1 –** Expression of genes (Z-scores) with an unadjusted *p* value of $p<0.0005$, in mice infected with *Heligmosomoides polygyrus* and administered proanthocyanidins (PAC), relative to infected mice (water-dosed controls).

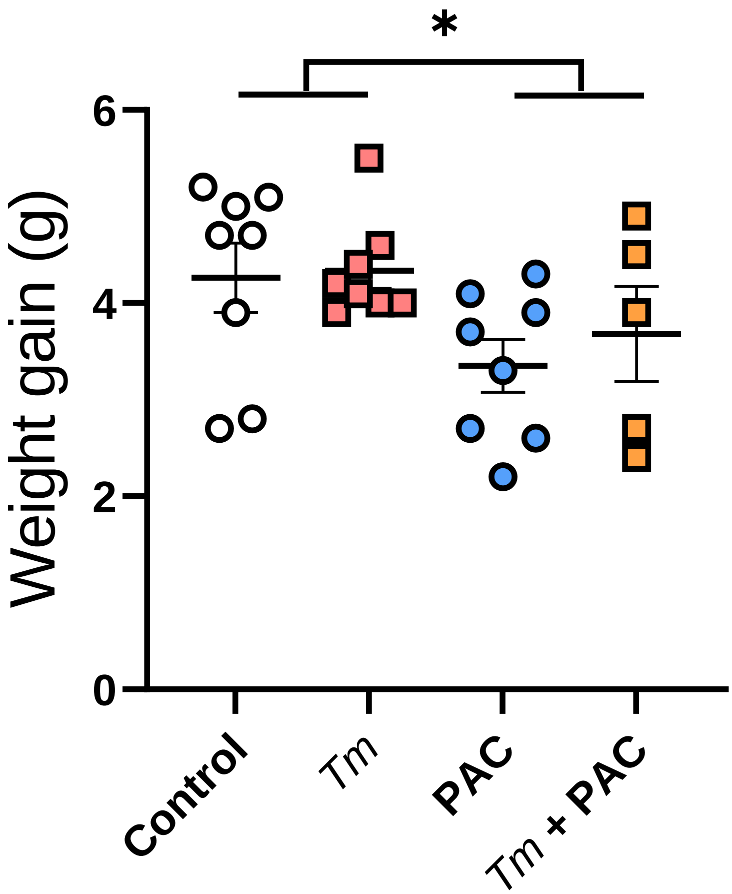


**Supplementary Figure 2 –** Weight gain of mice during 7 week experiment infected or not with *T. muris* (*Tm*), and given either proanthocyanidins (PAC) or water (control). * $p < 0.05$ by two-way ANOVA. Pooled data from two independent experiments.

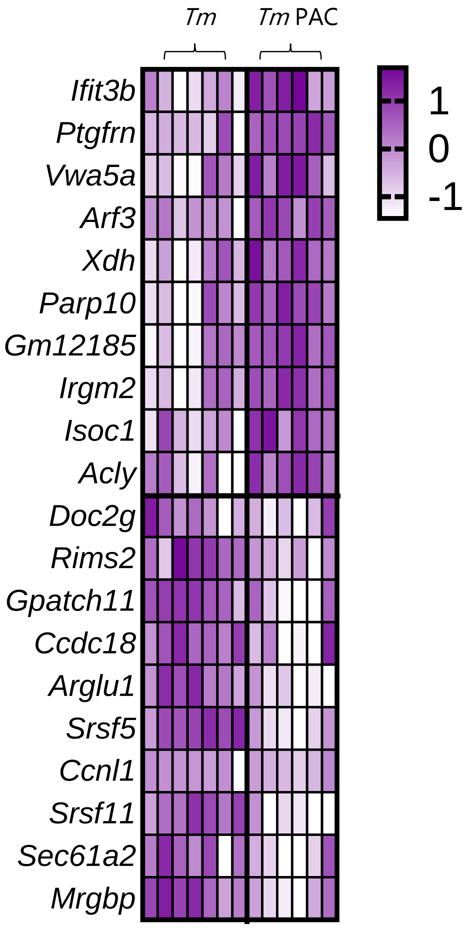


| Gene | padj | p value |
|---|---|---|
| Ifit3b | 3.00E-05 | 0.537435 |
| Ptgfrn | 0.001496 | 1 |
| Vwa5a | 0.001535 | 1 |
| Arf3 | 0.002261 | 1 |
| Xdh | 0.003576 | 1 |
| Parp10 | 0.00392 | 1 |
| Gm12185 | 0.004594 | 1 |
| Irgm2 | 0.004998 | 1 |
| Isoc1 | 0.005045 | 1 |
| Acly | 0.005209 | 1 |
| Doc2g | 0.002861 | 1 |
| Rims2 | 0.002063 | 1 |
| Gpatch11 | 0.001131 | 1 |
| Ccdc18 | 0.001098 | 1 |
| Arglu1 | 0.001011 | 1 |
| Srsf5 | 0.000817 | 1 |
| Ccnl1 | 0.000495 | 1 |
| Srsf11 | 0.000408 | 1 |
| Sec61a2 | 0.000265 | 1 |
| Mrgbp | 7.83E-05 | 0.702248 |

**Supplementary Figure 3 –** Expression of genes (Z-scores) with an unadjusted *p* value of $p<0.0005$, in mice infected with *Trichuris muris* and administered proanthocyanidins (PAC), relative to infected mice (water-dosed controls).

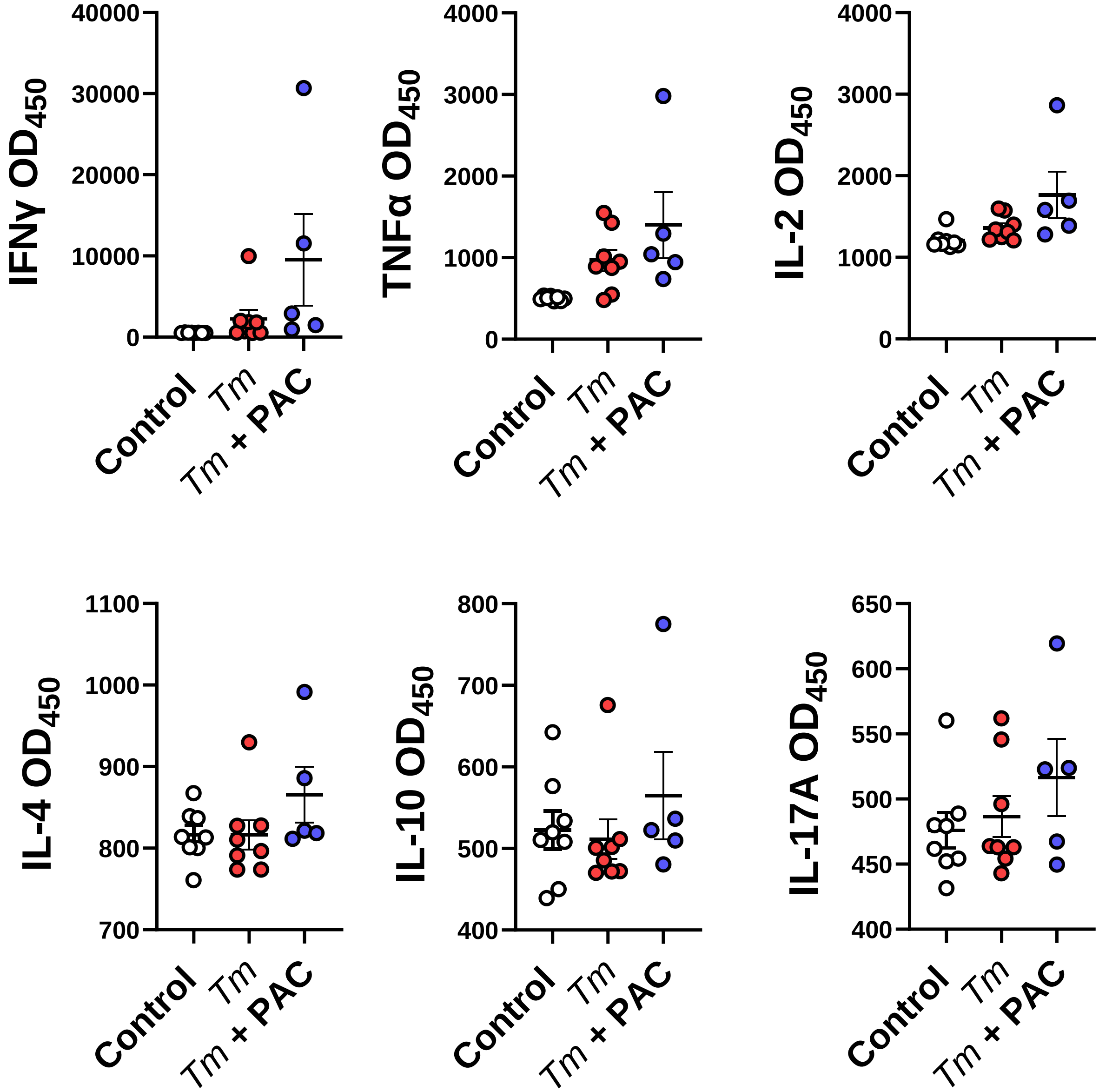


**Supplementary Figure 4 –** Cytokine production from mesenteric lymph node cells following stimulation with *Trichuris muris* antigen, in uninfected (control) mice, *T. muris* infected mice, and *T. muris* infected mice administered proanthocyanidins (PAC) or water (control). Pooled data from two independent experiments.

### *Hp* vs Control

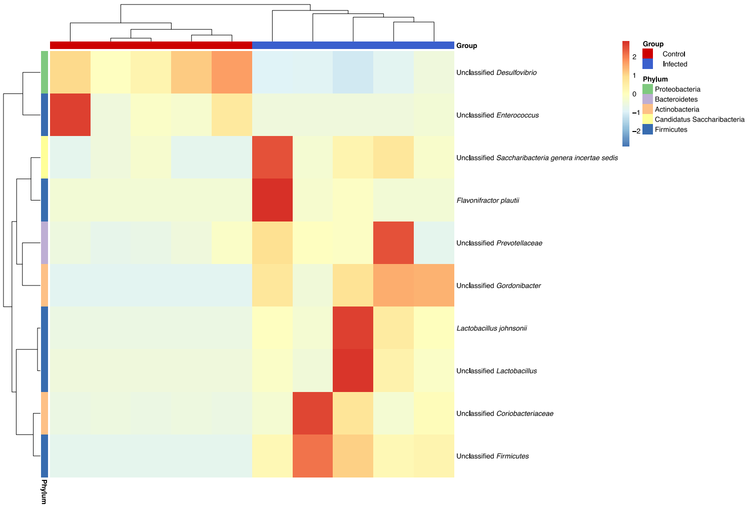


### *Hp* + PAC vs PAC

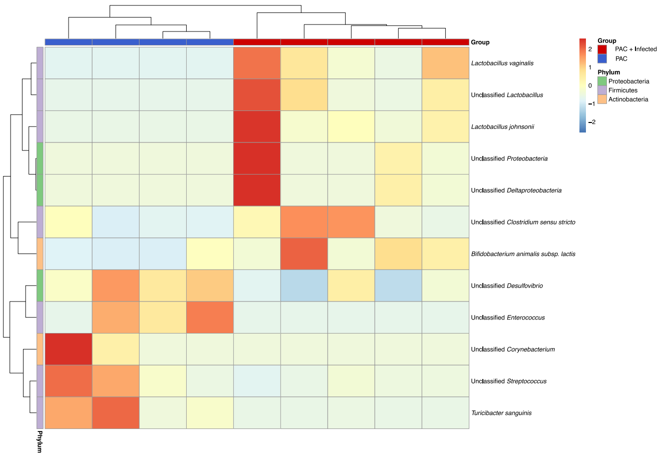


### PAC vs Control

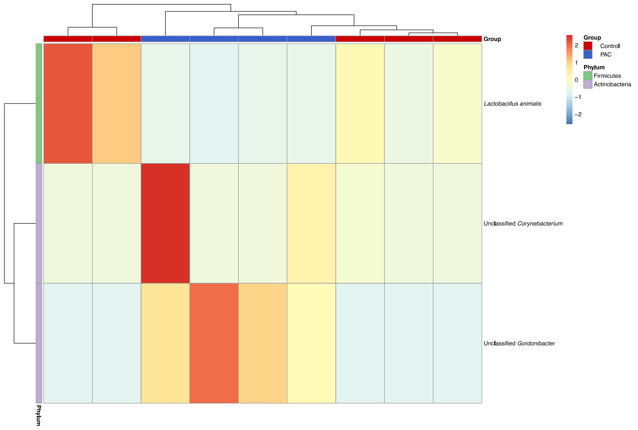


**Supplementary Figure 5 -** Species significantly impacted (adjusted *p* value <0.05) by *Heligmosomoides polygyrus* infection (*Hp*) and/or proanthocyanidin (PAC) intake

### *Tm* vs Control

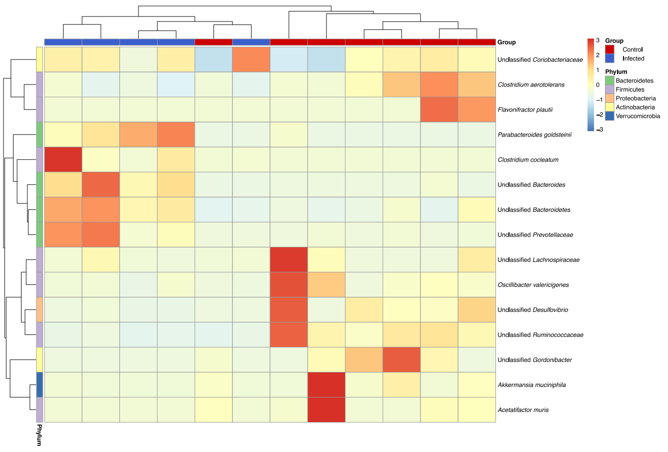


### *Tm* + PAC vs PAC

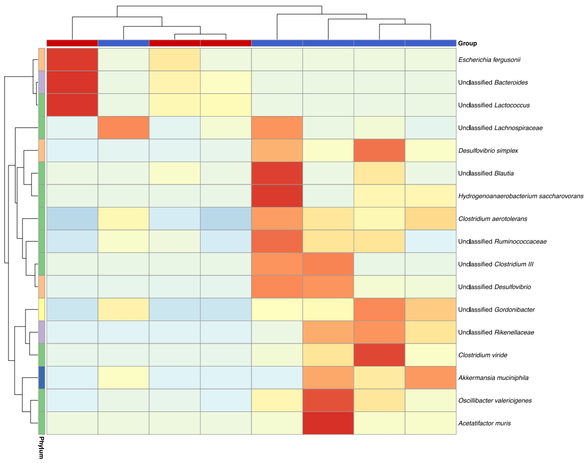


### PAC vs control

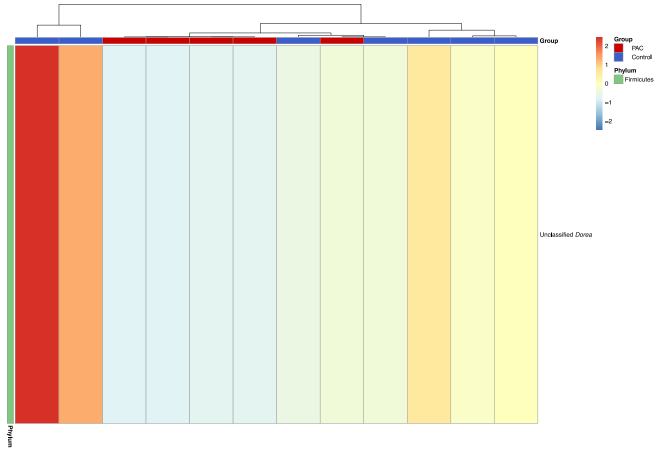


**Supplementary Figure 6** - Species significantly impacted (adjusted *p* value <0.05) by *Trichuris muris* infection (*Tm*) and/or proanthocyanidin (PAC) intake

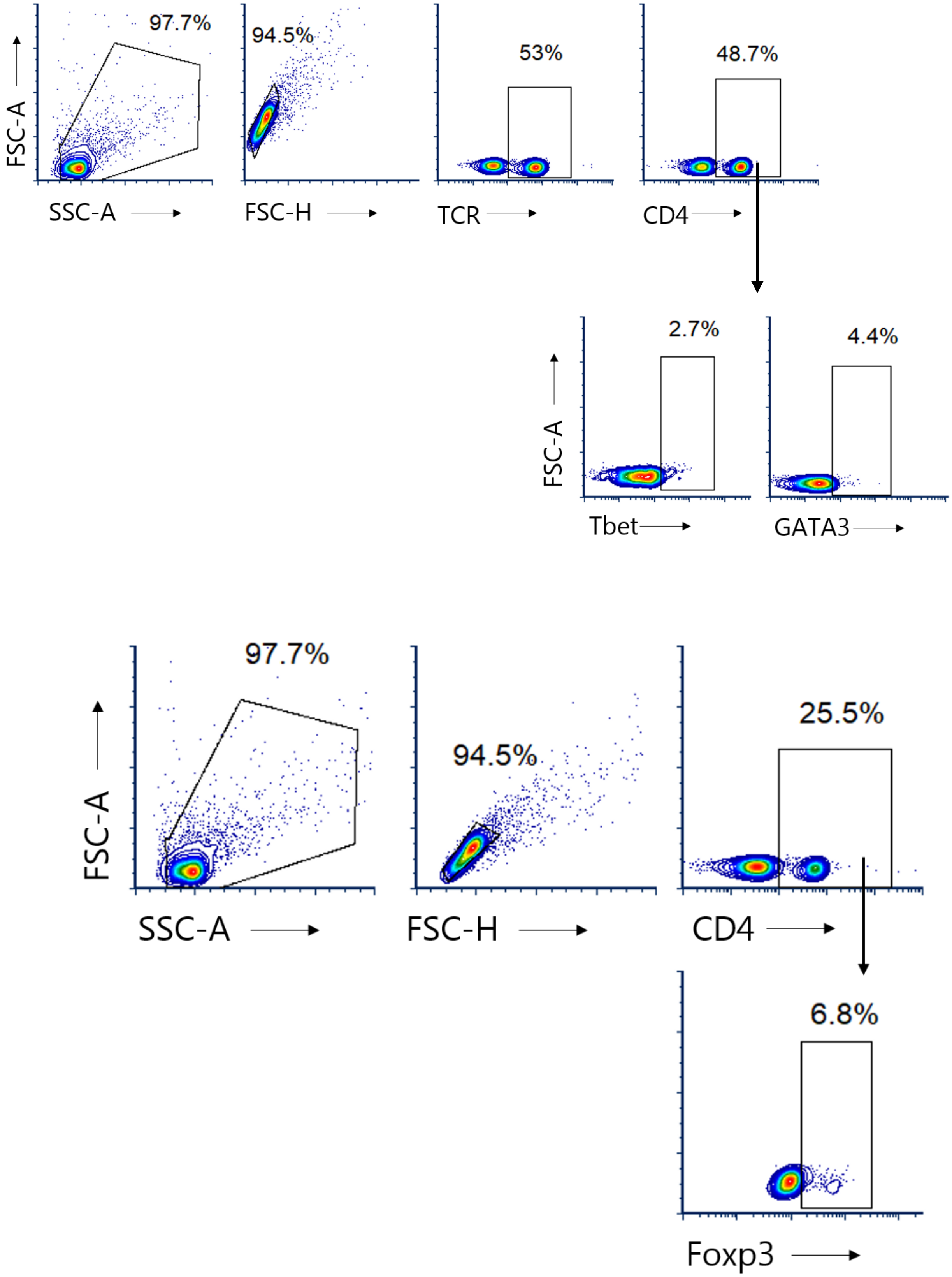


**Supplementary Figure 7 -** Gating strategy for mesenteric lymph model cell flow cytometry analysis. Cells were gated on live cells and doublets excluded. For Tbet and GATA3 staining, cells were gated on TCR and CD4 expression, followed by intracellular staining for the transcription factors. For Foxp3 staining, cells were gated on CD4 followed by intracellular staining for the transcription factor.

**File S1**

RNAseq (DESeq2) results comparing *H. polygyrus* infected mice to control mice

**File S2**

RNAseq (DESeq2) results comparing PAC-dosed mice to control mice (duodenum)

**File S3**

RNAseq (DESeq2) results comparing PAC-dosed, *H. polygyrus*-infected mice to *H. polygyrus*-infected mice

**File S4**

RNAseq (DESeq2) results comparing *T. muris* infected mice to control mice

**File S5**

RNAseq (DESeq2) results comparing PAC-dosed mice to control mice (caecum)

**File S6**

RNAseq (DESeq2) results comparing PAC-dosed, *T. muris*-infected mice to *T. muris*-infected mice
